# Spatial management can significantly reduce dFAD beachings in Indian and Atlantic Ocean tropical tuna purse seine fisheries

**DOI:** 10.1101/2020.11.03.366591

**Authors:** Taha Imzilen, Christophe Lett, Emmanuel Chassot, David M. Kaplan

**Affiliations:** Institut de Recherche pour le Développement (IRD), Avenue Jean Monnet, CS30171, 34203 Sète cedex, France; MARBEC, Univ Montpellier, CNRS, Ifremer, IRD, Sète, France; Sorbonne Université, Collège Doctoral, 75005 Paris, France; IRD, PO BOX 570, Victoria, Seychelles; Seychelles Fishing Authority, PO BOX 449, Victoria, Seychelles

**Author notes:** **Author email addresses:** (Taha Imzilen). **Corresponding author:** Taha Imzilen.

**Keywords:** Marine pollution, Fishing debris, Coral reefs, Fish aggregating device (FAD), Ocean currents

## Abstract

Debris from fisheries pose significant threats to coastal marine ecosystems worldwide. Tropical tuna purse seine fisheries contribute to this problem via the construction and deployment of thousands of man-made drifting fish aggregating devices (dFADs) annually, many of which end up beaching in coastal areas. Here, we analyzed approximately 40 000 dFAD trajectories in the Indian Ocean (IO) and 12 000 dFAD trajectories in the Atlantic Ocean (AO) deployed over the decade 2008-2017 to identify where and when beachings occur. We find that there is tremendous promise for reducing beaching events by prohibiting deployments in areas most likely to lead to a beaching. For example, our results indicate that around 40% of beachings can be prevented if deployments are prohibited in areas in the south of 8°S latitude, the Somali zone in winter, and the western Maldives in summer for the IO, and in an elongated strip of areas adjacent to the western African coast for the AO. In both oceans, the riskiest areas for beaching are not coincident with areas of high dFAD deployment activity, suggesting that these closures could be implemented with relatively minimal impact to fisheries. Furthermore, the existence of clear hotspots for beaching likelihood and the high rates of putative recovery of dFAD buoys by small-scale fishers in some areas suggests that early warning systems and dFAD recovery programs may be effective in areas that cannot be protected via closures if appropriate incentives can be provided to local partners for participating in these programs.

## 1 Introduction

Debris from fisheries pose significant threats to coastal marine ecosystems worldwide (Tavares et al. 2017; Parton et al. 2019). Tropical tuna purse seine fisheries contribute to this problem via their extensive use of drifting fish aggregating devices (dFADs) (Consoli et al. 2020). Whereas historically purse seine vessels divided there fishing effort between free-swimming fish schools and schools associated with naturally-occurring floating objects (FOBs), they increasingly focus principally on FOB fishing (Galland et al. 2016; Taconet et al. 2018). The attachment to FOBs of, first, radio beacons in the mid-1980’s and 1990’s and then satellite-tracked, GPS-equipped buoys from the early 2000’s, and most recently the integration of echo-sounders in satellite-tracked buoys have made this approach to catching tunas increasingly attractive to fishers (Chassot et al. 2014; Lopez et al. 2014). These technological developments have led purse seiners to manufacture and deploy large numbers of their own, manmade dFADs (Maufroy et al. 2017), and today it is believed that over 100 000 of these devices are deployed annually worldwide (Scott & Lopez 2014). dFADs typically consist of a floating structure and of a submerged substructure stretching up to 80 m below the surface (Imzilen et al. 2019). Some of the materials regularly used in dFAD construction include non-biodegradables such as PVC and metal tubes for the raft frames, ethylene vinyl acetate floats and plastic containers for buoyancy, and old nylon nets and pieces of salt bags for the subsurface structure. The massive increase in dFAD use poses a number of major concerns regarding ecological disturbance, overfishing, increased bycatch and creation of marine debris (Amandè et al. 2010; Dagorn et al. 2013; Filmalter et al. 2013; Maufroy et al. 2015). Most importantly for the context of this paper, a significant fraction of these dFADs end up beaching (i.e., stranding in coastal environments) (Maufroy et al. 2015), potentially damaging sensitive habitats such as coral reefs, and contributing to coastal marine debris and ghost fishing (Balderson & Martin 2015; Stelfox et al. 2016; Zudaire et al. 2018). This is of particular concern in a context of growing awareness of the extent of marine plastic pollution, with abandoned and lost fishing gears having been shown to be a major component of marine litter worldwide (Haward 2018; Lebreton et al. 2018; Richardson et al. 2019).

Given these concerns, dFAD beachings are a major area of interest for science, management and conservation. An initial examination of French dFAD spatiotemporal use in the tropical Indian Ocean (IO) and Atlantic Ocean (AO) over the period 2007-2011 indicated that ~10% of deployed dFADs ended up beached (Maufroy et al. 2015), highlighting the potential for considerable impacts on fragile coastal habitats due to these events. A similar examination in the Western and Central Pacific Ocean found that ~6% of all trajectories were likely to have beached over a two year period (2016-2017; Escalle et al. 2019). However, given the significant differences in bathymetry and circulation between the western and central Pacific Ocean, IO and AO, and the more than four-fold increase in the number of dFADs deployed by purse seiners in the IO and AO since 2011 (Katara et al. 2018; Floch et al. 2019), the extent to which existing literature applies to current patterns of dFAD use is an important open question. Moreover, the French fleet switched to almost exclusively using echo-sounder equipped dFAD tracking buoys around 2012 (Chassot et al. 2014; Floch et al. 2019) and other major purse seine fleets also started using this new technology on or before this date, potentially altering the spatio-temporal distribution of dFAD deployments, fishing activity and associated beaching events. In parallel, management measures have been taken by the tuna regional fisheries management organizations to limit the total number of GPS buoys used by each purse seine vessel in both the AO and IO, but these measures have not directly addressed the spatial and temporal dynamics of beachings and, therefore, their efficacy for reducing this problem is unknown. A new analysis of dFAD beachings focused on spatiotemporal patterns that might be useful for identifying appropriate mitigation measures to avoid beachings is therefore urgently needed.

The goal of this paper is to quantify the impacts of dFAD beachings and identify strategies for mitigating these impacts in the tropical IO and AO. Using a large dataset of over 50 000 dFAD buoy trajectories, we first extend and improve upon the analysis of Maufroy et al. (2015, 2018), estimating beachings for the decade 2008-2017 for the IO and AO. We then identify deployment locations likely to lead to beaching events, and, using this information, we are able to estimate the impact of closing high beaching risk areas to dFAD deployments on the overall beaching rate under a pair of reasonable fishing effort redeployment strategies. Results indicate that there is indeed much promise in the IO and AO for reducing dFAD beachings by implementing sensible spatial limitations on deployment locations.

## 2 Materials and methods

### 2.1 Data collection

Through a collaboration with the French frozen tuna producers’ organization (ORTHONGEL), the French Institute of Research on Development (IRD) has access to data on the locations of thousands of distinct GPS buoys attached to FOBs deployed by the French and associated flags (Mauritius, Italy, Seychelles) purse seine fleets operating in the tropical AO and IO from ~2007 onward (coverage ~75-86% before 2010 and ~100% after that date; Maufroy et al. 2015). Though GPS buoys can be attached to both natural FOBs and man-made dFADs, the vast majority of FOBs in both oceans are now man-made dFADs (>90% of buoy deployments in both oceans based on observer data for 2013-2017), and, therefore, we will refer to these buoy trajectories as dFAD trajectories even though a small fraction of them are for other types of objects. GPS buoys are attached to dFADs deployed at sea by purse seine fishing vessels and their associated support vessels. Buoys can also be exchanged on FOBs encountered at sea and the buoys retrieved from the water are generally brought back to port where they can be recovered by the owner vessel for reuse. A single GPS buoy may therefore be redeployed several times, potentially on different dFADs. It is therefore important to note that, in this paper, we use the term ‘dFAD’ to refer to the entire device consisting of the floating object itself and the attached GPS buoy, whereas, the term ‘buoy’ designates solely the GPS buoy.

Buoy location data are transmitted with a periodicity that varies along the buoy trajectory, generally ranging from 15 minutes to 2 days. Buoy positions were filtered to remove those that were emitted while the buoy was onboard using a Random Forest classification algorithm that is an improvement over that developed in Maufroy et al. (2015) (Appendix A). This improved classification algorithm is estimated to have an error rate of ~ 2% when predicting onboard positions and ~ 0.2% for at sea positions (Supplementary Table A4).

In this study, we used data of dFAD positions covering the decade 2008-2017. This data set consists of ~15 million IO positions representing a total of 38 845 distinct buoys and ~6 million AO positions representing a total of 12 147 distinct buoys. Separately, locations and times for dFAD deployments are available in French logbook data from 2013 onward.

### 2.2 Identification of dFAD beaching events

dFAD beachings were identified in two steps. The first step was to find dFADs that had an abnormally small rate of movement for an extended period of time, whereas the second step removed false positives (e.g., buoys onboard or at port) from this list of potential beachings. A given dFAD position was considered to be a potential beaching if: (1) at least 2 other later positions were within 200 m, and (2) all these close positions span a time period exceeding 1 day. The 200 m threshold is based on a dFAD snagged on the very bottom of its <100 m length nets hanging below the dFAD swinging at most 100 m in each direction. The time span of at least 1 day is required to avoid identifying as beachings multiple position emissions from a single buoy over a short time period, such as occurs when the emission periodicity of dFAD positions is modified to 15 min to facilitate detection by vessels before a fishing set.

In the second step, the putative beachings identified in this first step were filtered to remove non-beaching events based on 4 tests: (1) the beaching is more than 10 km from a major fishing port to avoid cases where dFAD buoys are at a port; (2) the beaching event is <5 km from land or the water column depth is <100 m; (3) all positions are classified at sea and there are no gaps in location emission exceeding 2 days over the 5 days preceding the beaching; (4) greater than 90% of all positions of a given buoy within the time span of the potential beaching event are associated with the beaching event (i.e., meet the distance criteria described above; this condition avoids cases where a buoy happens to pass multiple times through the same area, because of an eddy for example). Only beaching events meeting these 4 conditions were considered for further analyses.

About half of the beachings identified by the conditions described above occurred in the water. The other half were generally located on land close to small fishing ports or coastal villages (Supplementary Fig. B2, Fig. B3 and Fig. B4). This suggests that these buoys were retrieved by small-scale boats, likely fishers. As these boats generally intercept dFADs in coastal areas and only collect the buoy for its valuable electronics, leaving the raft and netting to drift, it is entirely possible that these dFADs (without the buoy) later ended up beaching. Nevertheless, given the uncertainty regarding the fate of these dFADs, calculations in this paper have been carried out both including all beachings and including only beachings in the water. Unless otherwise stated, statistics reported in the paper are for all beachings including those on land. In the following sections of this paper, beachings located in water and on land are respectively referred to as “beachings along shore” and “recoveries displaced to shore”.

The number of beaching events per km of the continental shelf was calculated by counting all beachings occurring in each 5°x5° grid cell and then dividing that number by the kilometers of continental shelf edge, defined by the 200 m isobath, within the cell. The continental shelf edge was used instead of the coastline to avoid anomalously high beaching rates for some very small islands surrounded by large continental shelf areas.

For identifying beachings, classifying beachings as on land or at sea and determining the continental shelf edge, coastline data were obtained from OpenStreetMap land polygons (available at https://osmdata.openstreetmap.de/data/land-polygons.html; accessed 2020-02-19) and bathymetry was obtained from the 30-arcsecond-resolution General Bathymetric Chart of the Oceans (GEBCO v.2014; available at https://www.gebco.net/data_and_products/gridded_bathymetry_data; accessed 2020-02-19).

### 2.3 Drift locations leading to beachings

In order to identify dFAD drift locations that had a high risk to lead to a beaching event, we calculated the fraction of buoys that beach within 3 months of a passage through a given 1°x1° grid cell. This analysis was carried out over the entire study period, but also by season to estimate seasonal variability in beaching risk. We selected 3 months as the time limit as it is intermediate between the mean timespan of at sea trajectories and that of the lifespan of a buoy in the dataset (i.e. 25 and 196 days, respectively), and because 3 months was considered a reasonable timespan over which fishers and managers could reasonably be expected to predict and mitigate for beaching likelihood. To ensure that results are not strongly sensitive to this choice, additional analyses were carried out to calculate the fraction of buoys that beach within 12 months. Note that individual buoy trajectories were separated into multiple in water trajectories using breaks defined by gaps of more than 2 days or positions classified as onboard representing more than 1 minute of trajectory time. The 1 minute limit was imposed to remove very short trajectory segments that were problematic for the classification algorithm (Appendix A).

Since beachings threaten fragile marine habitats, especially coral habitats, we carried out the same analyses focusing exclusively on beachings in coral reef areas. Data on the global distribution of coral reefs were obtained from UNEP-WCMC, WorldFish Centre, WRI, TNC (2018, version 4.0; available at https://data.unep-wcmc.org/datasets/1; accessed April 30, 2019).

### 2.4 Deployment risk

To assess potential for reducing the dFAD beaching rate, we investigated closing areas of high beaching risk to dFAD deployments. Deployment locations were obtained from logbook data, whereas proportion of beaching was estimated as described above. Logbook deployment data was used instead of putative deployments from reconstructed dFAD trajectories because, though the random forest position classification model has a very high accuracy rate and predicted deployment locations do follow the spatial distribution of logbook deployment locations (Maufroy et al. 2015), accurately predicting deployment locations is quite difficult and error prone given that a single error anywhere in the trajectory will split the trajectory, generating a new false deployment (Maufroy et al. 2015). Given the high quality of logbook data, it was considered that this was the most accurate estimate of recent dFAD deployment locations.

Multiplying dFAD deployments by the proportion of devices beaching allowed us to predict the reduction in beachings that would result from closing a given area. Different size areas corresponding to specific percentages of all pre-closure deployments were closed in order of beaching risk going from highest to lowest. Two hypotheses were considered regarding the number and spatial distribution of deployments after closing an area to deployments: (1) closures eliminate deployments that would have occurred in closed areas (i.e., fishing effort reduction occurs), and (2) closures displace deployments formerly in closed areas to remaining unclosed areas in proportion to the relative density of deployments prior to implementation of closures (i.e., “fishery squeeze” occurs; Halpern et al. 2004).

## 3 Results

The number of French buoys deployed per year has increased dramatically and continuously over the decade 2008-2017, especially in the Indian Ocean (Fig. 1a). Over that period, more GPS buoys were deployed in the Indian Ocean (~ 40 000) than in the Atlantic Ocean (~ 12 000). The percentage of all deployed dFADs that ended up beaching has also dramatically increased from ~3.5% in 2008 to ~20% in 2013 (Fig. 1b; these numbers are roughly halved if we count only beachings along shore). After 2013, the percentage of dFADs that beached stabilizes at ~15-20% in the IO and ~19-22% in the AO. In total, we obtained 7187 beaching events for the IO and 2283 for the AO.

**Fig. 1.**
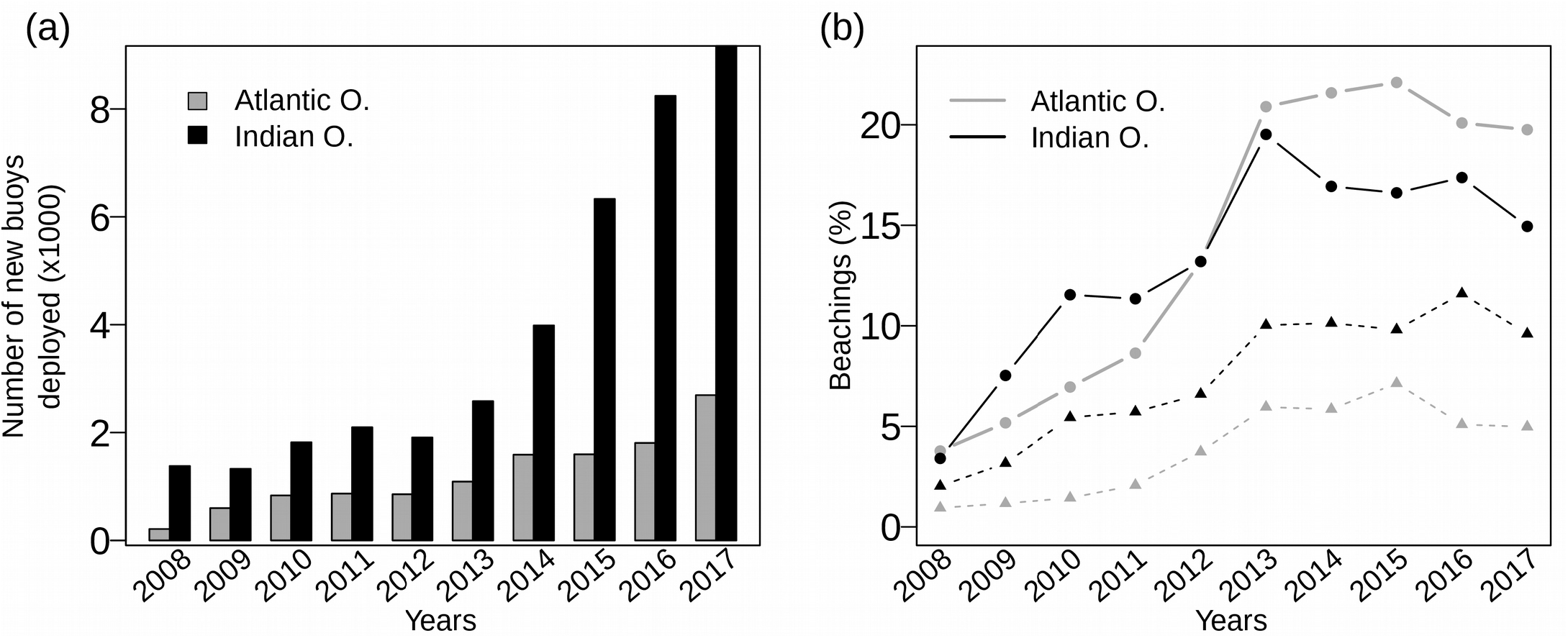
(a) Annual number of new buoys deployed by the French and associated flags purse seine fleet in the Atlantic (grey) and Indian (black) oceans over the period 2008-2017 and (b) percentage of these buoys that beached. The lines in (b) with solid circles include all beachings, whereas the lines with solid triangles include only beachings identified along shore. Beachings along shore and recoveries displaced to shore were separated via intersection with OpenStreetMap land polygons.

Maps of these 9470 beaching locations clearly identify coastal beaching hotspots (Fig. 2a and Supplementary Fig. C1a). Beachings occur in several zones in the IO, including southern Somalia, Kenya, Tanzania, Seychelles and the Maldives. In the AO, they occur mainly along the west African coast and the Gulf of Guinea between 20°N and 20°S. In both oceans, beachings also sporadically occur in more remote areas outside typical purse-seine fishing grounds (Maufroy et al. 2017), such as Indonesia, South Africa, Brazil and the Caribbean. Including only beachings that occur along the shore, the number of beaching decreases mostly along the western and north-eastern African coasts and in the Maldives (Fig. 2b and Supplementary Fig. C1b), which indicates that significant rates of putative recovery of dFAD buoys occur in those areas.

**Fig. 2.**
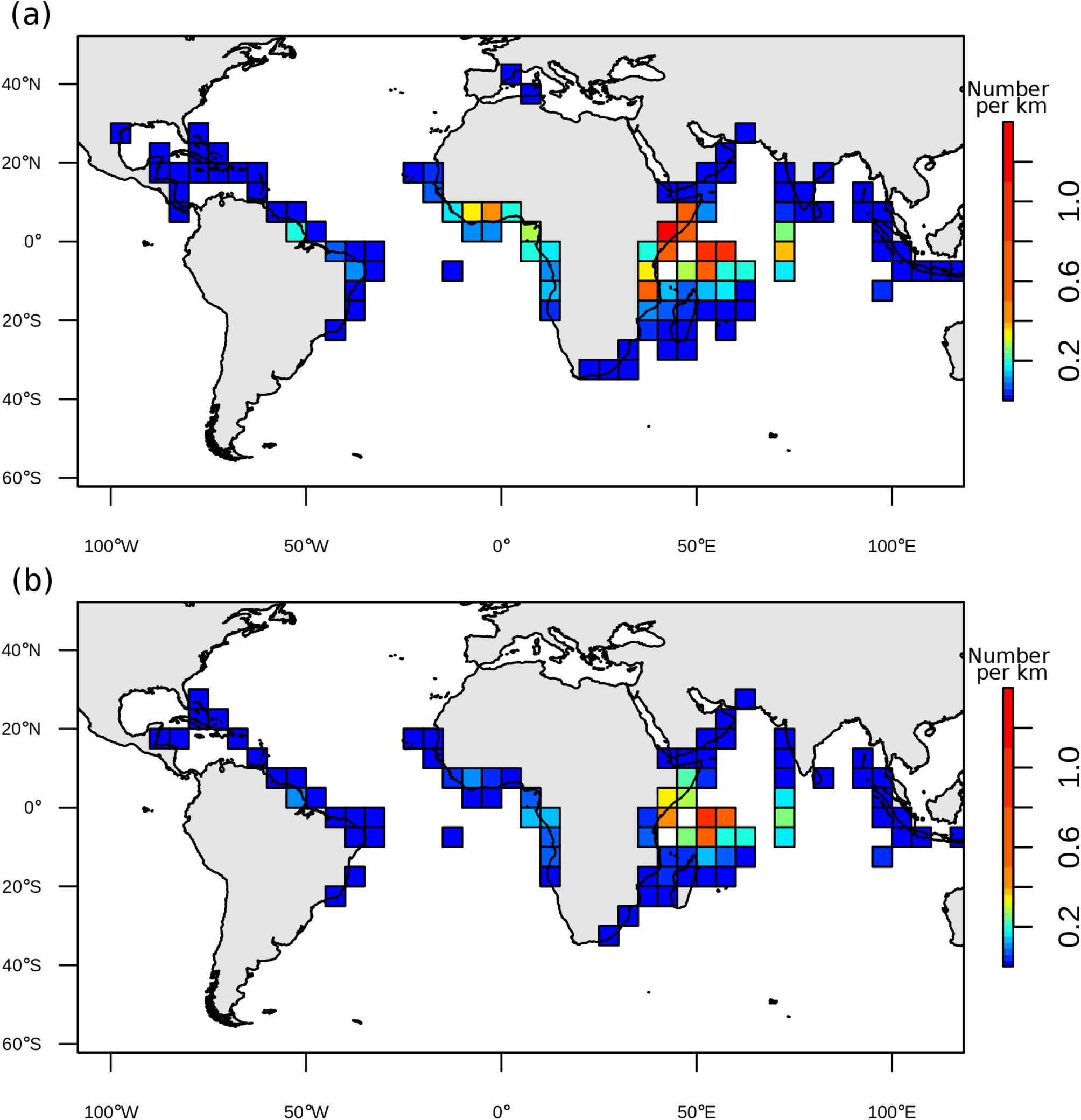
The number of French dFAD beachings recorded in our data per km of continental shelf edge in each 5°x5° grid cell for the period 2008-2017. Darker areas indicate higher rates of beaching. In (a), all beachings are considered, whereas in (b) only beachings along shore are included. Beachings along shore and recoveries displaced to shore were separated via intersection with OpenStreetMap land polygons. Note that our dFAD trajectory data is incomplete before ~2010, so the absolute number of beachings per kilometer is likely somewhat higher than values shown in the figure, though differences are likely to be small as the number of dFADs was far lower before 2010 than after 2010.

In both oceans, the proportion of dFADs beaching within 3 months of passing through a 1°x1° grid cell shows high spatial heterogeneity, with hotspots of beaching likelihood clearly visible (Fig. 3a). In the IO, the Gulf of Aden, Oman, Mozambique Channel, eastern and northern Madagascar, northern Maldives, western India, Sri Lanka and western Indonesia are all high risk areas for beaching. In the AO, the Gulf of Guinea, southern West Africa, the northern coast of South America and Caribbean have high proportion of beaching. Including only beachings that occur along shore reduces beaching proportions in all areas and reduces the importance of some coastal areas characterized by a high density of small-scale fishers, such as in the vicinity of the Arabian Peninsula, the northern Gulf of Guinea and West Africa (Fig. 3b). Increasing the temporal window from 3 months to 12 months increases somewhat the spatial area over which proportion of beaching is non-negligible, but overall spatial patterns remain the same (Supplementary Fig. C2). Seasonal variability in dominant currents impacts beaching risk in predictable ways. For example, in the IO, during the winter monsoon, onshore currents create an area of high proportion of beaching east of Somalia, but this high risk area disappears during the upwelling favorable period of the summer monsoon (Supplementary Fig. C4). However, seasonal variability in the AO was weak. Finally, focusing exclusively on dFAD beachings on coral reefs narrowed the areas of high beaching risk to the north-west of the Maldives, Seychelles, northern Madagascar, the Mozambique channel and the Caribbean (Supplementary Fig. C5).

**Fig. 3.**
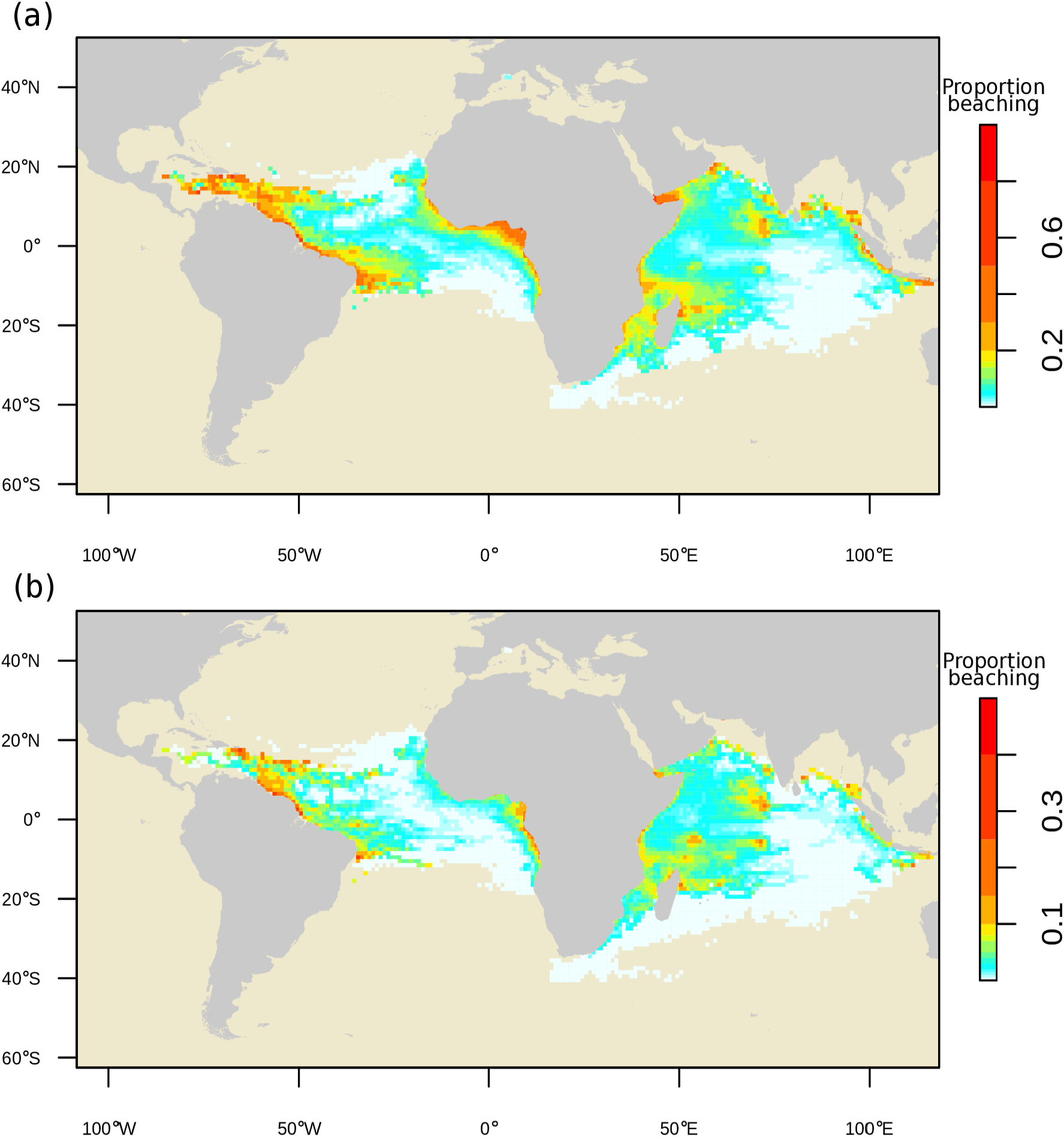
Maps of the proportion of dFADs that beached within 3 months after passing through each 1°x1° grid cell over the period 2008-2017. In (a), all beachings are considered, whereas in (b) only beachings along shore are included. Beachings along shore and recoveries displaced to shore were separated via intersection with OpenStreetMap land polygons. Note that the color intervals are unevenly distributed to highlight the low values.

Major areas of dFAD deployments during 2013-2017 spanned the whole fishing grounds of the French and associated flags purse seine fishery (Fig. 4a-b). In the AO, dFADs were deployed all along the coast of West Africa, from Mauritania down to Angola with the the most intense activity being observed along the equator and off the coasts of Mauritania, Gabon and Angola. In the IO, dFADs were deployed in the Western Indian Ocean, including the Exclusive Economic Zones of the Seychelles, Comoros, Kenya, French overseas territories and northwest of Madagascar in the northern Mozambique Channel. dFADs deployments were particularly frequent North-West of the Seychelles.

**Fig. 4.**
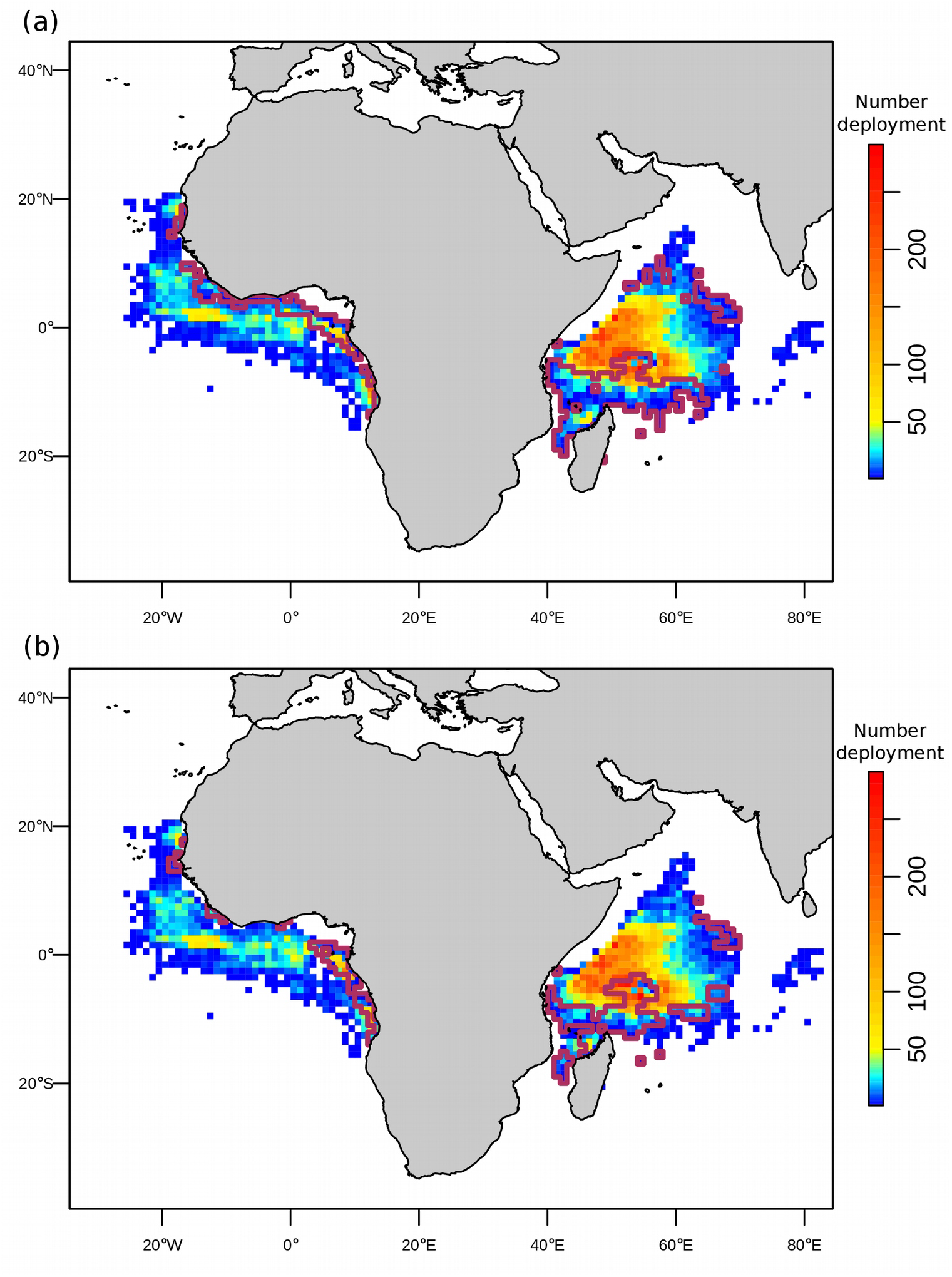
Density maps representing the number of dFAD deployments in each 1°x1° cell recorded in logbook data for the period 2013-2017. The thick, solid curves delimit areas representing the 20% of deployments most likely to produce a beaching within 3 months of a dFAD passing through those areas. In (a), all beachings are considered, whereas in (b), only beachings along shore are included. Beachings along shore and recoveries displaced to shore were separated via intersection with OpenStreetMap land polygons.

Combining spatial proportions of dFADs that beached (Fig. 3a-b) with observed dFAD deployment positions (Fig. 4a-b), we estimated the expected change in beachings and dFAD deployments due to prohibiting dFAD deployments in the highest risk areas for both oceans. Under all scenarios of dFAD deployment redistribution, spatial prohibitions are predicted to significantly reduce beaching rates. For example, if we prohibit dFAD deployments in areas corresponding to the 20% of deployments with highest beaching risk, we can prevent 37% of beachings in the IO and 40% in the AO in the absence of dFAD deployment effort redistribution, and 21% and 25% of beachings in the IO and AO, respectively, even if we allow for dFAD deployment redistribution to areas with less beaching risk (Fig. 5a). These percentages are even higher when we focus on the proportion of beaching including only beachings that happen along shore, with up to a 52% reduction in beachings in the AO even if the total number of deployments is conserved via effort redistribution (grey dashed line in Fig. 5b).

**Fig. 5.**
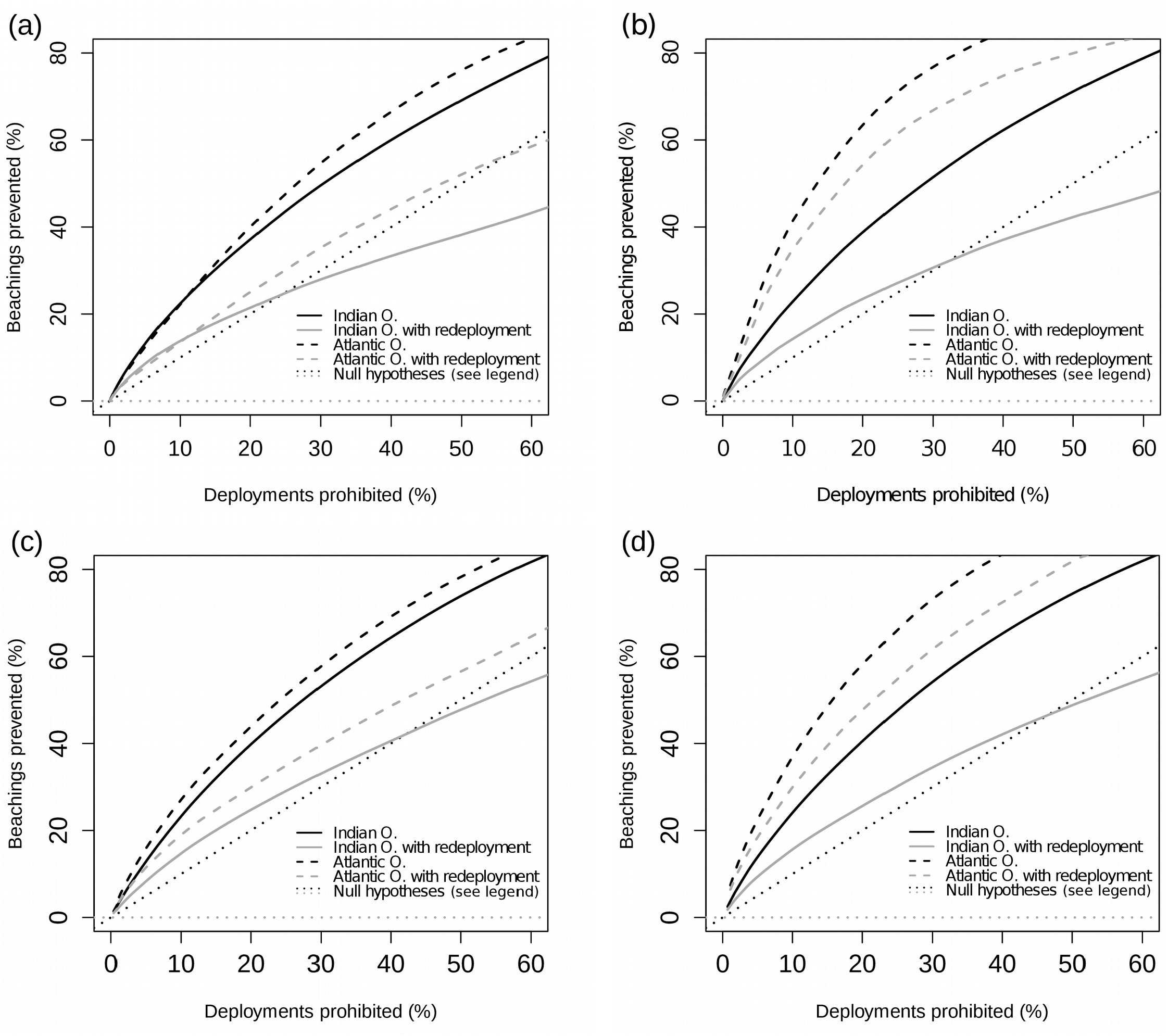
Predicted reduction in beaching rate as a function of the amount of area put aside in annual (a-b) or quarterly (c-d) closures to dFAD deployments. Areas are closed from most likely to least likely to produce a beaching within 3 months of deployment, with area being quantified along the x-axis in terms of the fraction of deployments that occurred in closed areas prior to their closure. Black and grey dotted lines correspond to the null expectation of what the corresponding black and grey curves would look like if all areas had the same beaching risk, and are the same in the IO and AO. In (a) and (c), all beachings are considered, whereas in (b) and (d), only beachings occurring along shore are included. Beachings along shore and recoveries displaced to shore were separated via intersection with OpenStreetMap land polygons.

Spatial prohibitions can be optimized to account for seasonal variability in beaching risk. For example, if areas corresponding to the 20% of deployments in areas with the highest beaching risk for each quarter are closed to dFAD deployments (Supplementary Fig. C6), we predict a 27% and 28% reduction in the IO and AO, respectively, even if dFAD deployment redistribution is allowed (Fig. 5c).

Focusing exclusively on beachings in coral reefs, prohibiting the 20% of deployments in the IO with the highest beaching risk to corals reduces coral reef beachings by 27% assuming dFAD deployment redistribution (Supplementary Fig. C7b), but the zones prohibited differ significantly from those that would be prohibited to reduce all beaching events (compare Fig. 4a and Supplementary Fig. C7a).

Closing the highest beaching risk areas to dFAD deployments is particularly effective at reducing beaching events in the south-western IO and in the eastern Gulf of Guinea in the AO (Fig. 6). If one focuses exclusively on coral reef beaching, then significant beaching reductions in the IO are also seen in the Maldives and off Indonesia (Supplementary Fig. C8). These results apply to both with and without dFAD deployment redistribution scenarios.

**Fig. 6.**
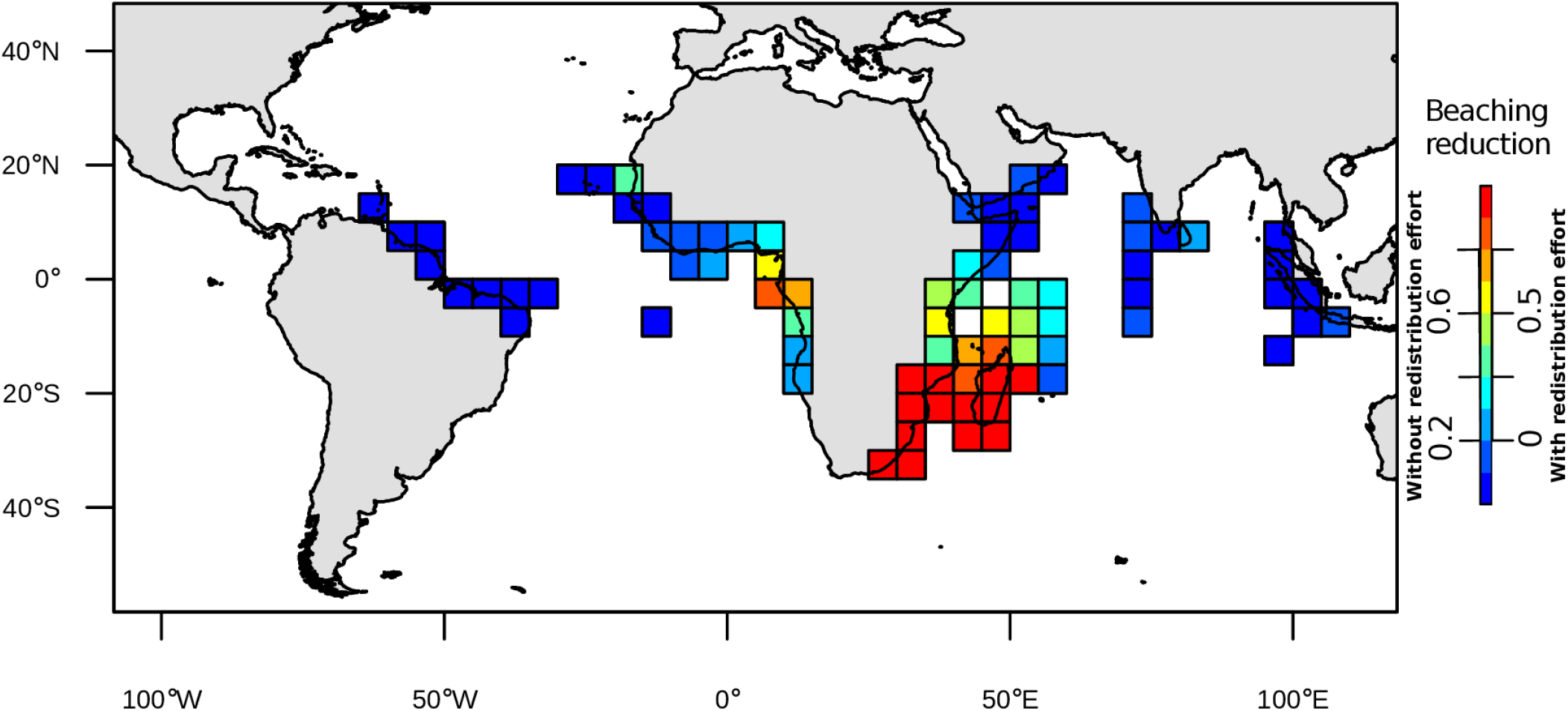
Map representing the predicted reduction in beaching when the 20% of dFAD deployments most likely to produce a beaching within 3 months are prohibited (see areas in Fig 4a), without (values on the left of the colorbar) and with (values on the right of the colorbar) dFAD deployment effort redistribution to non-prohibited areas.

## 4 Discussion

The overriding conclusion to be drawn from our results is that there is potentially a lot to be gained in terms of reduction in the rate of dFAD beachings from spatio-temporal closures for dFAD deployments by purse seine fishing vessels in the IO and AO. We examined a wide range of scenarios for closure objectives, implementation, and post-closure effects: considering all beachings versus just strandings along shore; considering all coastal zones versus just coral reefs; implementing static versus quarterly varying closures; and post-closure effort reduction versus effort redistribution to remaining open areas. In all cases, closing the riskiest areas to beaching is predicted to produce a tremendous reduction in beachings. Analyses of recent dFAD deployments in the IO by the Spanish fleet (the dominant other fleet in both oceans) indicate that Spanish and French deployments have quite similar spatial distributions. This suggests that our results may be applicable to all fleets (Katara et al. 2018), though access to dFAD trajectory data should be enhanced to confirm this. Perhaps most encouraging, high risk areas generally are relatively coherent in space so that it should be feasible from a management perspective to implement closures (e.g., south of 8°S in the IO and coastal zones in the Gulf of Guinea in the AO). In both oceans, the riskiest areas for beaching are not coincident with areas of high dFAD deployment activity nor fishing activities (Maufroy et al. 2015), suggesting that these closures could be implemented with relatively minimal impact to fisheries. The beaching reduction across coastal areas spared by the closures for dFAD deployment is highest in the south-western IO and in the eastern Gulf of Guinea in the AO, suggesting that our proposed deployment closure strategy is particularly efficient to protect these areas. The north-western IO and the northern Gulf of Guinea, which both represent hotspots of beaching, are less protected by the closures for dFAD deployments. However, high rates of putative recovery of dFAD buoys by coastal boats in these areas indicate that beaching early warning systems and dFAD recovery programs may be effective in areas that cannot be protected via closures if appropriate incentives can be provided to local partners for participating in these programs

As reported elsewhere (Maufroy et al. 2015; Floch et al. 2017, 2019), the number of dFADs deployed in both oceans has dramatically increased over the last decade. More surprising, the fraction of dFADs that end up beaching increased significantly over the period 2008-2013, after which time the fraction stabilizes. As this 2008-2013 period is coincident with a number of changes in the fishery, such as the switch to echosounder buoys (2010-2012), an increase in the prevalence of dFAD fishing as opposed to fishing on free-swimming schools (Assan et al. 2019; Floch et al. 2019) and the fallout from Somali piracy (~2007-2011), it is hard to assign a specific cause to this pattern. One hypothesis is that as the number of dFADs has increased, the fraction of dFADs that are never fished upon has become more and more important to the point that after 2013 the fraction beaching simply reflects the balance one would expect in the absence of fishing between dFADs that beach versus dFADs that sink at sea. The stabilization of the beaching rate after 2013 may also be partially due to the implementation after 2014 of industry and/or regional fisheries management organizations limit on the number of buoys monitored by purse seine vessels (ICCAT 2019; IOTC 2019a) as fishers may remotely deactivate non-productive dFADs to remain under industry limits, resulting in the loss of location information for these FADs that continue to drift at sea and may later beach.

The risk of beaching depends strongly on the upper ocean circulation and its seasonal variability. In the IO, the southern African coast represents a high beaching risk area throughout the year due to the westward flowing Northern Equatorial Madagascar Current (Schott et al. 2009) that drives dFADs to the coasts of Mozambique and Tanzania. In the northern IO, high beaching risk areas change with monsoon regimes. The Somali coast represents a high beaching risk area in the winter when the Somali Current flows westwards (Schott & McCreary 2001), but not during the summer, when the western Maldives become a high risk area due to monsoon driven eastward circulation. There is less effect of seasonality on beaching risk in the AO, where areas of high beaching risk are driven by more-stable dominant circulation patterns. Along the western coast of Africa, beachings are related to the North Equatorial Countercurrent and the Guinea Current flowing eastwards, whereas high risk areas along the northern coast of South America and the Caribbean are linked to the South Equatorial, North Equatorial, North Brazil and Caribbean Currents flowing westwards (Bourles et al. 1999).

Our estimates of dFAD beaching rates after 2013 are higher than those estimated in the western central Pacific (Escalle et al. 2019) and in previous examinations in the IO and AO (Maufroy et al. 2015; Zudaire et al. 2018). Escalle et al. (2019) examined an area of the Pacific characterized principally by many small island chains, perhaps explaining lower beaching rates with respect to the continental land masses of the IO and AO. In the IO and AO, Maufroy et al. (2015) examined the period prior to 2013 for which we also find lower beaching rates. Zudaire et al. (2018) were principally concerned with the more-limited area of the Seychelles Archipelago, which is composed of a large set of small islands similar to the area examined by Escalle et al. (2019) in the western central Pacific, and they considered a somewhat more restrictive definition of beaching.

There have been several recent management changes regarding the use of dFADs that may alter future dFAD beaching patterns, highlighting the importance of continuous monitoring of dFAD trajectories. The Indian Ocean Tuna Commission (IOTC) and the International Commission for the Conservation of Atlantic Tunas (ICCAT) currently limit the number of buoys monitored by an individual purse seine vessel at any given time to 300 (ICCAT 2019) and 350 (IOTC 2019a) buoys in the AO and IO, respectively, and these limits are likely to decrease over time. The IOTC has also implemented a resolution to progressively reduce and phase out the number of support vessels that assist the purse seiners with the management of dFADs (IOTC 2019b). These changes may lead purse seine vessels to optimize their use of dFADs in a number of ways. One potential outcome would be that fishers remotely deactivate dFADs that are likely to beach or drift outside of areas of interest so as to remain under industry limits. This practice is of much concern as it would result in the loss of information on the extent and location of dFAD beachings currently made available via fishing companies on a voluntary basis. Tuna regional fisheries management organizations should put in place appropriate incentives or other measures to assure that this information loss does not occur.

This study would not have been possible without access to a long and extensive time series of data on French dFAD trajectories. Though access to these extensive datasets is still quite limited for most fishing fleets worldwide, there are a number of encouraging signs of increased reporting of dFAD deployments and other dFAD activities to tuna regional fisheries management organizations (IOTC 2019a). We are hopeful that comprehensive datasets from all purse seine fishing fleets will be available in the near future, permitting better estimates of the impacts of management options and the development of real-time tools for the management of dFAD impacts on marine ecosystems.

## 5 Supporting information

Supporting information available online comprises details of the new classification model for onboard and at sea states of dFAD trajectory data (Appendix A), and quantification of beachings occurred in water (beachings along shore) and on land (recoveries displaced) (Appendix B), as well as additional figures presenting the number of French dFADs beached in each 5°x5° cell, proportions of beaching using a 12 month time window, seasonal variability in beaching risks and beaching risks for coral reefs (Appendix C).

## Acknowledgments

This work was funded by the Research Project INNOV-FAD (European Maritime and Fisheries Fund, measure n°39, OSIRIS #PFEA390017FA1000004, and France Filière Pêche), the European Research Project CECOFAD2 (Specific Contract No 9 of EASME/EMFF/2016/008) and the Ob7 Exploited Tropical Pelagic Ecosystems Observatory of the IRD. We express our sincere thanks to ORTHONGEL for making their dFAD tracking data available and to the Ob7 for data management and preparation. We are particularly grateful to L. Dagorn, D. Gaertner, A. Maufroy, L. Floch and M. Goujon for their assistance. We also thank two anonymous reviewers for their useful comments.

