## Supporting information for "Spatial management can significantly reduce dFAD beachings in Indian and Atlantic Ocean tropical tuna purse seine fisheries"

#### Contents:

**Appendix A - New Classification model for onboard and at sea states of dFAD trajectory data**

Provides details of the new random forest classification model for onboard and at sea states of dFAD trajectory data used to extract at sea trajectories from dFAD tacking buoy data.

**Appendix B - Quantification of beachings as in water (beachings along shore) or on land (recoveries displaced)**

Details of the methodology used to identify, separate and classify "beachings" occurring in water (beachings along shore) from those occurring on land (recoveries displaced).

**Appendix C - Additional figures**

Additional figures presenting the number of French dFADs beached in each 5°x5° cell, beaching probabilities using a 12 month time window, seasonal variability in beaching probabilities and beaching probabilities for coral reefs.

**Appendix D - Interactive map showing the locations of recent individual dFAD beachings**

Presentation of a website with an interactive map showing the locations of recent individual dFAD beachings.

### Appendix A: New Classification model for onboard and at sea states of dFAD trajectory data

**4 Problem of positions with very short time steps** **3**

**References** **3**

#### 1 Overview

French dFAD trajectory data contain a mixture of geographic positions that were emitted while the transmitting buoy was onboard a boat and while the buoy was in the water. The onboard positions must be eliminated to recover the at sea trajectories useful for identifying beaching events. We accomplished this using a random forest classification algorithm. The algorithm used here is an improvement over that presented by Maufroy et al. (2015). In particular, it is distinguished with respect to the previous algorithm by:

1) The inclusion of additional training data derived from more recent dFAD trajectories
2) The use of new predictor variables based on variability in buoy speed or temperature for a set of positions immediately temporally before and after the position to be classified

#### 2 New training dataset

The original classification algorithm developed by Maufroy et al. (2015) was based on a training dataset consisting of 204 buoys and 18,357 classified positions (Table A1). Data was manually classified based on buoy speed and temperature and data on nearby fishing activity. In this original dataset, classified buoys were randomly drawn from the period 2009-2010.

Since the time that this training dataset was developed, the number and type of French dFAD buoys has considerably changed. In particular, the buoy manufacturer is now Marine Instruments. These buoys contain echosounders and have a higher maximum emission rate than previous models (<15 minutes versus 1 hour or more in previous models). As these technological changes potentially impact both classification success and dFAD deployment and use, we decided to include additional training data drawn from more recent time periods in our classification model.

New training data was classified manually with the aid of a Shiny application in R that permitted simultaneous visualization of buoy speed, temperature, direction, emission rate and echosounder information, as well as dFAD deployment, visit and recovery data drawn from recent fisher logbooks and observer data. This additional training data consisted of 172 buoys and 61,419 classified positions drawn from the time period 2012-2018 (Table A2). Though classified buoy trajectories were largely randomly drawn from the set of buoy trajectories that possessed both logbook and observer data, 10-20 trajectories were specifically selected because they possessed data in the vicinity of Sri Lanka during periods when the buoy had been recovered by artisanal fishers and was transported to a Sri Lankan port. This data was known to be difficult to properly classify due to the low speed of artisanal fishing boats, and additional classification data from this area was found to visually improve classification success in this region.

### 3 New classification model

In addition to including both new and old training data, the classification algorithm we used included a different set of predictor variables. Some predictor variables that were found to have little predictive power by Maufroy et al. (2015), such as rate of change in temperature, were removed, and new variables based on a temporal window around the position to be classified were added.

#### 3.1 New predictor variables

New predictor variables included:

- the amount of time before and after a given position that buoy speed was less than 3 m/s
- the standard deviation of speed and temperature in a 7 position window around a given position (3 before, 3 after and the given position)

The first of these was included because speeds greater than 3 m/s were extremely rare in water positions, but common in boat positions (Fig. A1), so even though an individual boat position may have a low speed, subsequent or preceding positions were likely to have high speed (Fig. A2). Similarly, both speed and temperature were observed to be more variable when buoys were onboard, and therefore the standard deviation of these variables in a small temporal window around a position to be classified was found to be effective at separating onboard and at sea positions (Figs. A3 & A4).

The full set of predictor variables is presented in Table A3.

#### 3.2 Model description

The final classification model as executed in R is presented below.

```
randomForest(formula = class ~ lt_5km_land + dist_port_km + mean_time_change_s +
    mean_speed_ms + abs_azimuth_change_180 + acceleration_ms2 +
    local_speed_ms_stddev + local_water_temp_stddev + time_lt_3ms_s +
    before_time_lt_3ms_s + after_time_lt_3ms_s + is_mi_buoy,
    data = themodel$training.data, ntree = 1500, mtry = 4)
```

The meanings of the different predictor variables are given in Table A3.

#### 3.3 Model diagnostics

Not surprisingly, the classification model has prefect prediction success when applied to the training data used to calibrate the model (Table A4; note, however, that this was not the case for the model used in Maufroy et al. 2015, which had imperfect prediction success even when applied to the training data used to construct the model). However, internal cross validation in the random forest model suggests that there is an ~2.3% error rate when predicting onboard positions and an ~0.2% error rate when predicting at sea positions (Table A5). These error rates are considerably lower than those estimated by Maufroy et al. (2015), which predicted a mean error rate over both position classes (i.e., onboard and at sea) of ~2.2%. The superior performance of the new model was further confirmed by cross-application of the new model to the old training data and the old model to the new training data, which indicated at least a 50% reduction in error rate with the new classification model.

Model internal error rate indicates that the number of trees used is more than sufficient to reach model predictive stability (Fig. A5). By far the most important predictor variables for prediction success were the standard deviation of speed in a 7-position window around a given position (i.e., *local_speed_ms_stddev*), total time around a position for which buoy speed was less than 3 m/s (i.e., *time_lt_3ms_s*) and the speed of the buoy at a given position (i.e., *mean_speed_ms*) (Fig. A6). Though other variables contributed to model estimations, their impact was far weaker.

In Maufroy et al. (2015), a post-processing step was used to reclassify isolated pairs of consecutive onboard ($B$) positions as at sea ($W$) positions, which improved the overall classification success. Tests indicated that this post-processing step produced no significant improvement for the new classification model, likely due to the inclusion of new predictor variables that already take into account the temporal context of a given position. We, therefore, decided not to use this post-processing step with the new classification model.

#### 4 Problem of positions with very short time steps

A small number of buoys in the dFAD trajectory database possessed a small number of positions that were less than 60 seconds (most often less than 5 seconds) after the position immediately preceding it (3,387 buoys out of 64,613 buoys in the entire database). These are generally associated with recent buoys transmitting multiple sets of echosounder data, one set immediately after the other (e.g., due to multiple echosounder frequencies). The buoy speeds associated with these very small time steps were often anomalously large as they were presumably dominated by the noise in GPS position acquisition, which would be large relative to the small size of the time step. Given the large and rapidly varying velocities associated with these positions, they were often erroneously classified as on board a vessel. To correct for this, time periods less than 1 minute that were classified as on a vessel were ignored when dividing trajectories into at sea and onboard portions.

| Position classification | Number of buoys | Number of positions | Percent of positions |
|---|---|---|---|
| B | 197 | 2677 | 14.6 |
| W | 201 | 15680 | 85.4 |
| **Total** | **204** | **18357** | **100.0** |

Table A2: Number of buoys and positions by class in new training dataset. Position classification $B$ indicates onboard (i.e., boat) positions, whereas classification $W$ indicates at sea (i.e., water) positions. Note that the 'Number of buoys' column indicates the number of classified buoys that possess at least one position of a given class, and, therefore, an individual buoy can be counted in both the $B$ and $W$ classes.

| Position classification | Number of buoys | Number of positions | Percent of positions |
|---|---|---|---|
| B | 147 | 4893 | 8 |
| W | 167 | 56526 | 92 |
| **Total** | **172** | **61419** | **100** |

Table A3: Names and descriptions of the predictor variables used in the classification model.

| Predictor variable | Description |
|---|---|
| lt_5km_land | A boolean (true/false) indicating whether or not the buoy is within 5 km of the coast as determined by the GSHHS high resolution coastline data (http://www.soest.hawaii.edu/pwessel/gshhg/) |
| dist_port_km | Distance in kilometers to the nearest tuna fishing port |
| mean_time_change_s | Mean of the time step in seconds for the trajectory segment proceding a given buoy position and for the trajectory segement after the position |
| mean_speed_ms | Mean of the buoy speed in m/s over the trajectory segment proceding a given position and over the trajectory segement after the position |
| abs_azimuth_change_180 | Heading change in degrees between the trajectory segments proceding and immediately after the position |
| acceleration_ms2 | The linear acceleration (i.e., change in direction is *not* accounted for) of the buoy over the trajectory segements immediately proceding and after the position |
| local_speed_ms_stddev | The standard deviation of the buoy speeds in m/s over the trajectory segments immediately proceeding a set of 7 buoy positions in a window centered on the given position (3 before, 3 after and the central position) |
| local_water_temp_stddev | The standard deviation of the water temperature measurements in degrees Celcius at 7 buoy positions in a window centered on the given position (3 before, 3 after and the central position) |
| before_time_lt_3ms_s | The time period in seconds proceding a given position that the buoy speed was consistently less than 3 m/s |
| after_time_lt_3ms_s | The time period in seconds after a given position that the buoy speed was consistently less than 3 m/s |
| time_lt_3ms_s | The total time period in seconds before and after a given position that the buoy speed was consistently less than 3 m/s |

| Predictor variable | Description |
|---|---|
| is_mi_buoy | A boolean (true/false) indicating if the buoy in question was manufactured by Marine Instruments or not. Marine Instruments data corresponds to newer buoys typically including an echosounder |

Table A4: Confusion matrix for the new classification model when predicting on the combined training dataset used to construct the model. Position classification $B$ indicates onboard (i.e., boat) positions, whereas classification $W$ indicates at sea (i.e., water) positions.

| | Predicted B | Predicted W |
|---|---|---|
| **Observed B** | 7,570 | 0 |
| **Observed W** | 0 | 72,206 |

Table A5: Internal confusion matrix for new classification model. This confusion matrix is the result of internal cross-validation of the random forest model. Position classification $B$ indicates onboard (i.e., boat) positions, whereas classification $W$ indicates at sea (i.e., water) positions.

| | Predicted B | Predicted W | Classification error rate |
|---|---|---|---|
| **Observed B** | 7,393 | 177 | 0.0234 |
| **Observed W** | 158 | 72,048 | 0.0022 |

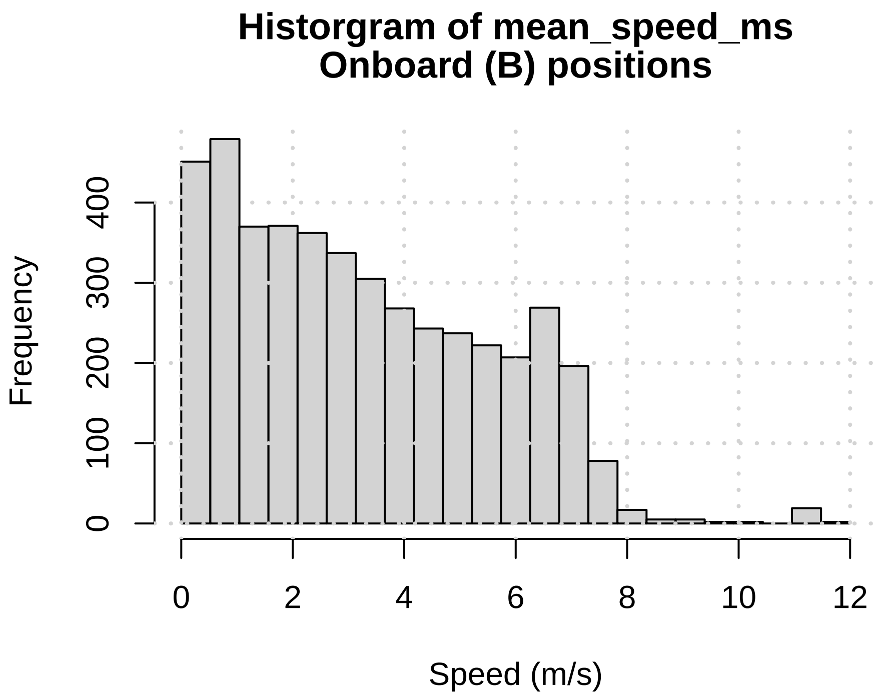

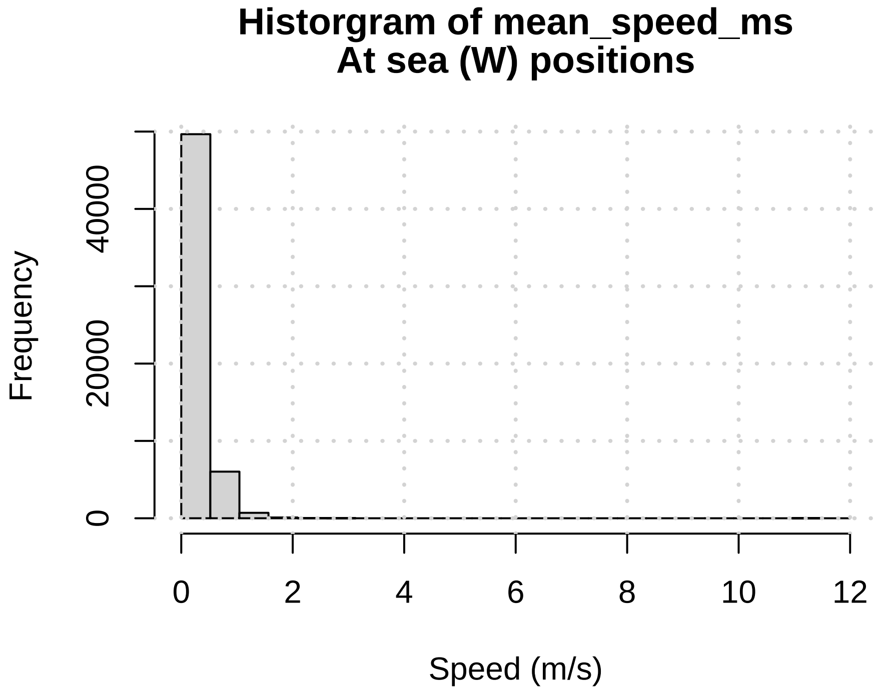

Figure A1: Histogram of *mean_speed_ms*, i.e., the mean of the two buoy speeds over the trajectory segments immediatiely proceding and immediately after a given buoy position (see Table A3 for more details). The top panel is for positions classified as onboard, whereas the bottom panel is for at sea positions.

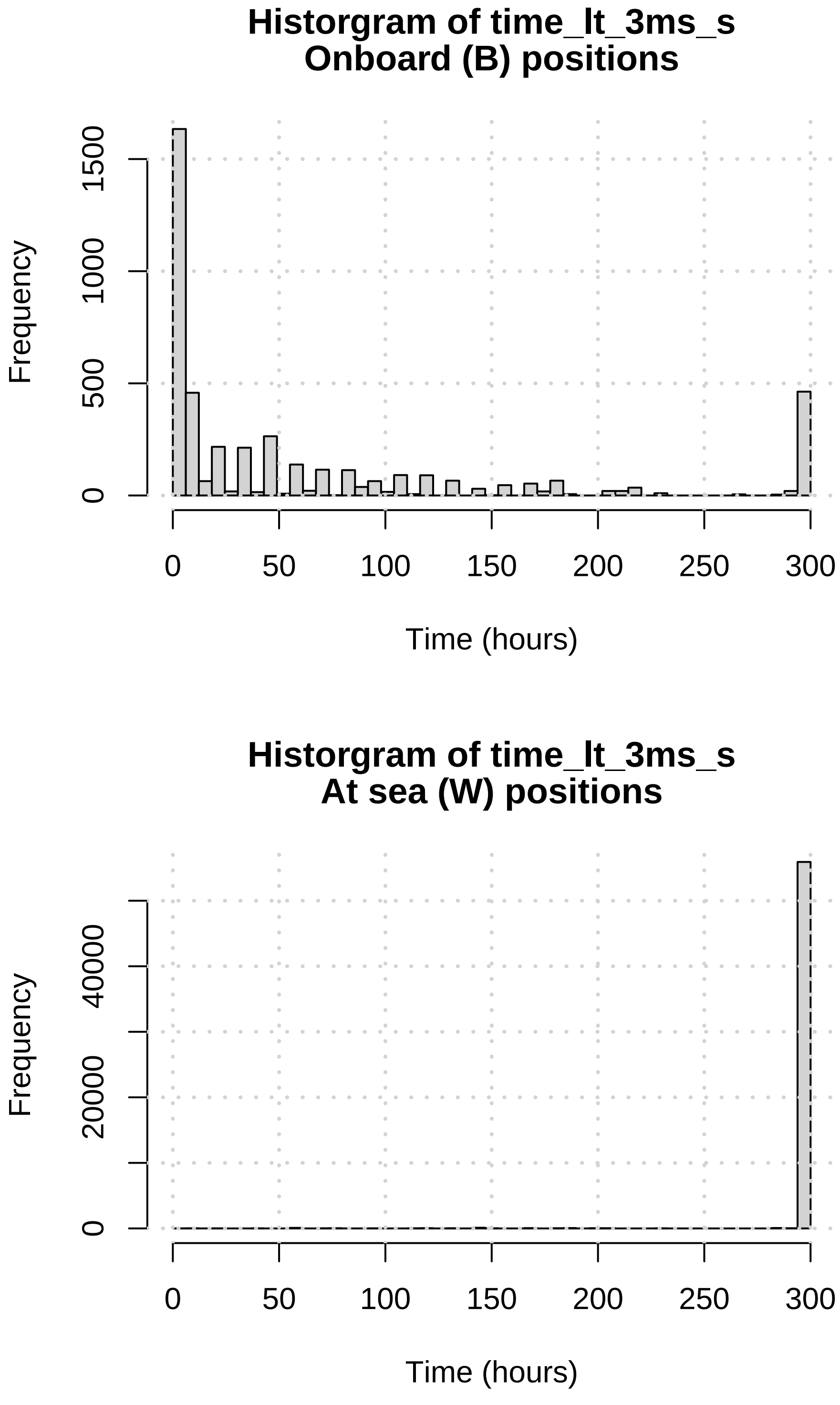

Figure A2: Histogram of *time_lt_3ms_s*, i.e., the time period around a given position for which buoy speed was inferior to 3 m/s (see Table A3 for more details). The top panel is for positions classified as onboard, whereas the bottom panel is for at sea positions. Values have been divided by 3600 to present results in units of hours instead of seconds.

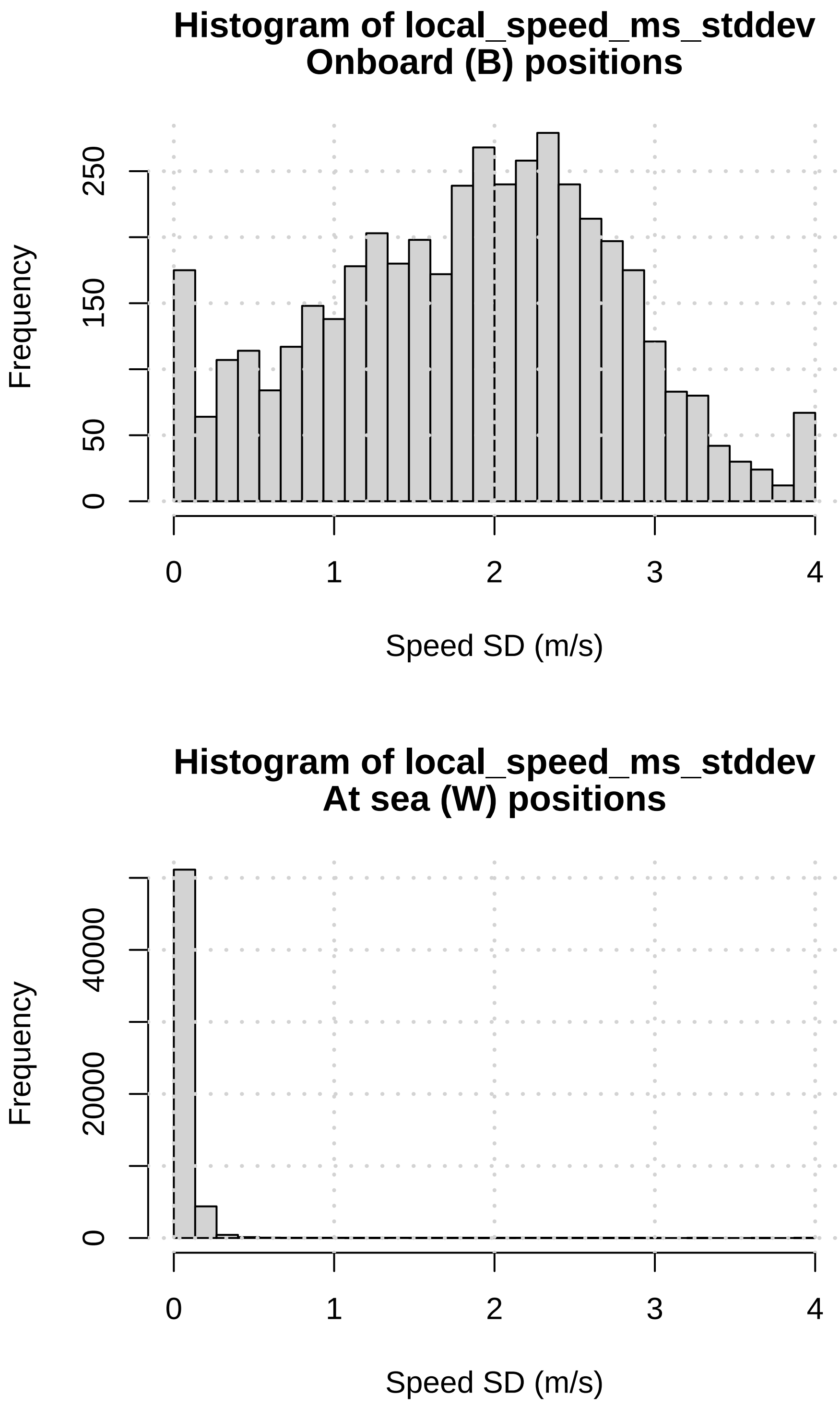

Figure A3: Histograms of *local_speed_ms_stddev*, i.e., the standard deviation of the buoy speed for a set of 7 consecutive buoy positions centered around a given position (see Table A3 for more details). The top panel is for positions classified as onboard, whereas the bottom is for at sea positions.

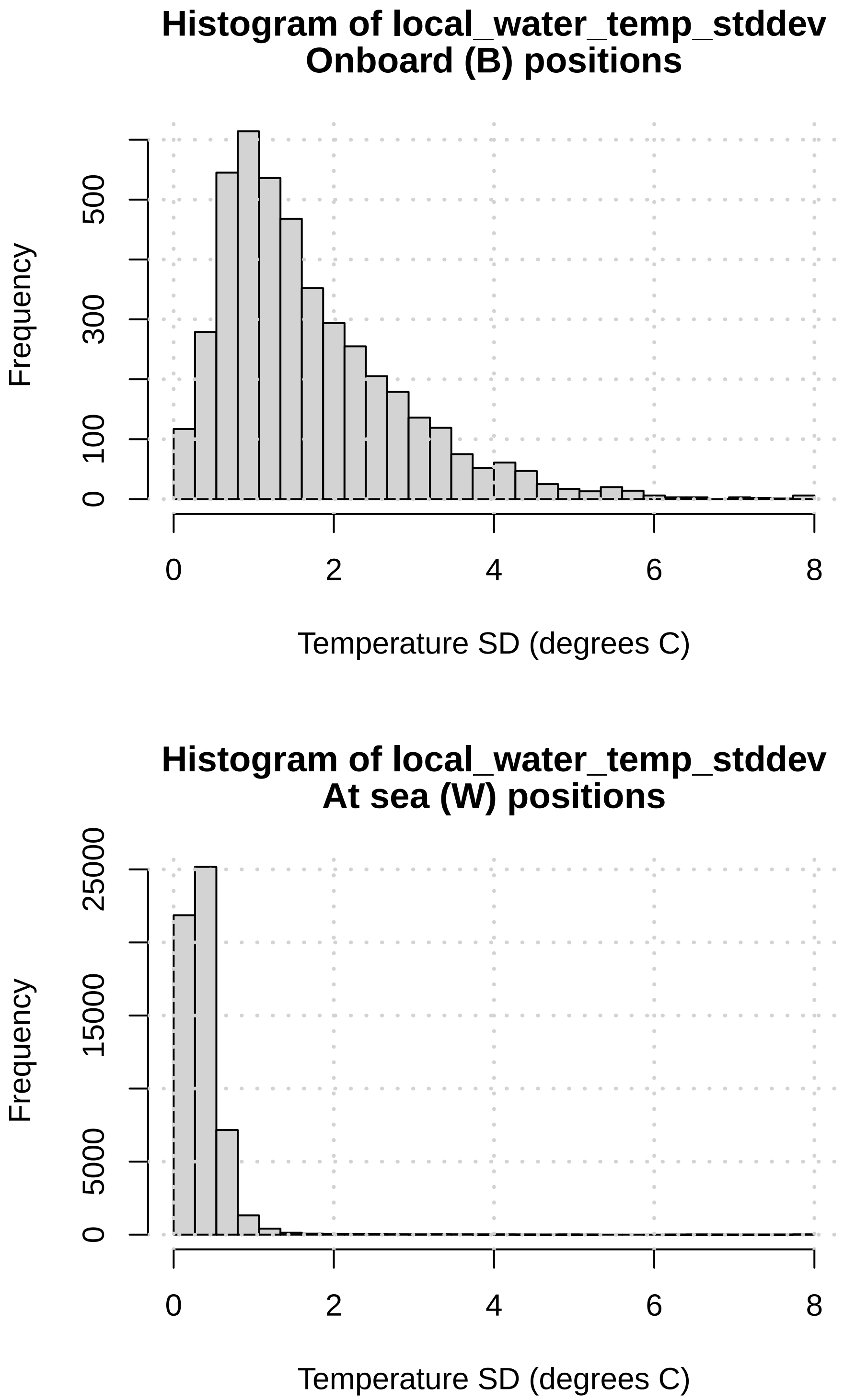

Figure A4: Histograms of *local_water_temp_stddev*, i.e., the standard deviation of water temperature measurements for a set of 7 consecutive buoy positions centered around a given position (see Table A3 for more details). The top panel is for positions classified as onboard, whereas the bottom is for at sea positions.

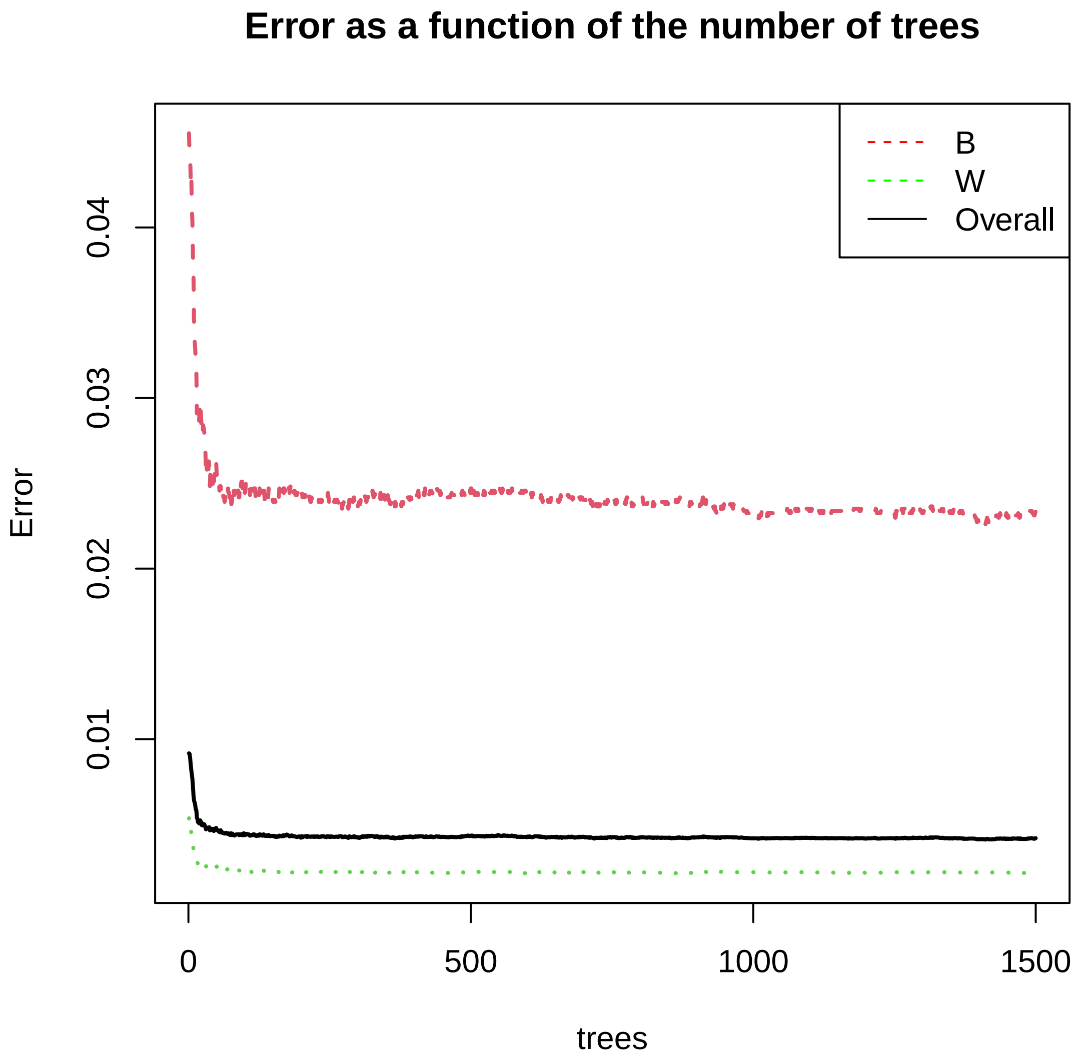

Figure A5: Error rate as a function of the number of trees included in the random forest model. Position classification $B$ indicates onboard (i.e., boat) positions, whereas classification $W$ indicates at sea (i.e., water) positions.

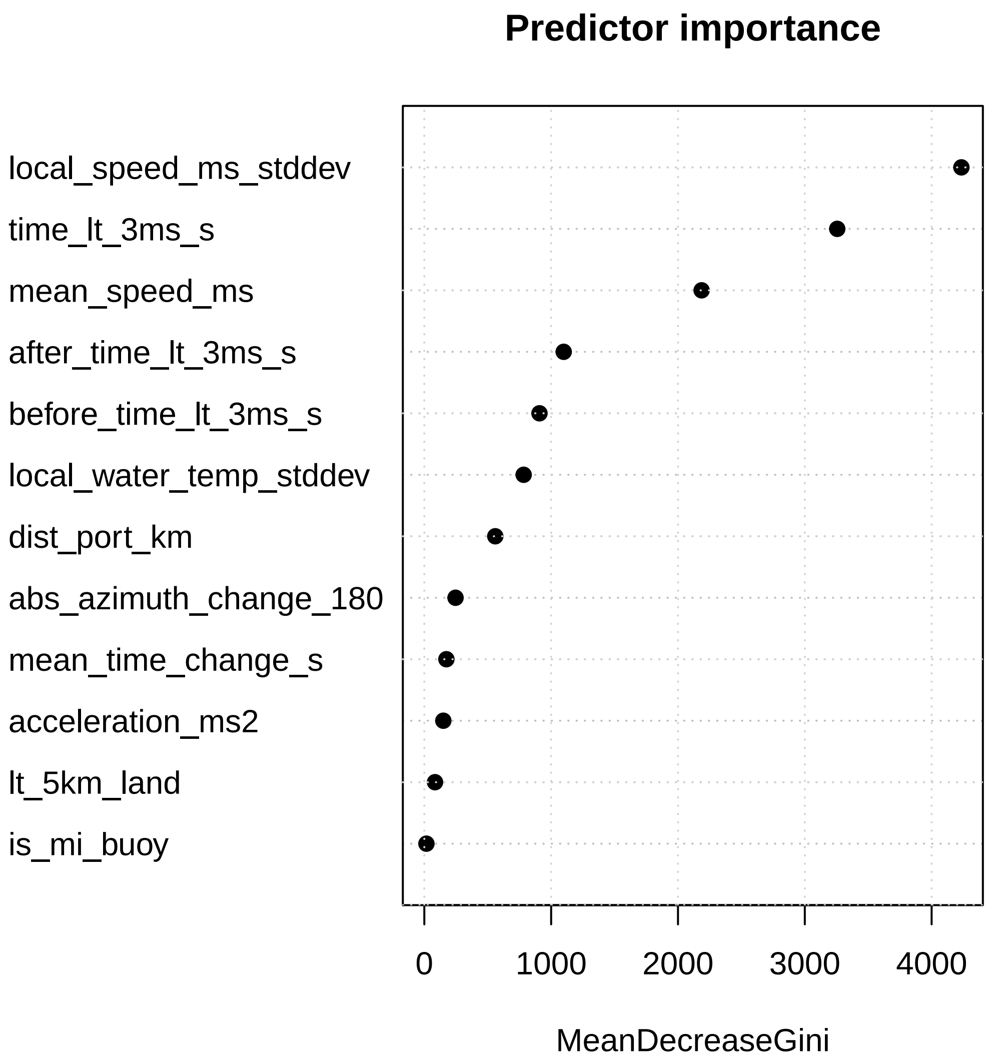

Figure A6: Predictor variable importance in new random forest classification model. See Table A3 for the meaning of the different explanatory variables.

### Appendix B - Quantification of beachings as in water (beachings along shore) or on land (recoveries displaced)

In a preliminary analysis of the beaching locations identified in this study, we found that most beachings occurred in water, stranding close to the coast, as expected. However, we also found an unexpectedly large amount of beaching locations on land, mostly within small ports or coastal villages. To further explore that critical aspect of our work we first classified manually a randomly selected sample of 100 identified beaching locations based on a Google Map visualization and obtained 55 beachings occurring in water and 45 on land. Moreover, beachings identified in water showed two situations: (i) beachings that occurred in water only (35 cases, e.g., Fig. B1) and (ii) beachings that occurred in water first and then found beached again on land some days later (20 cases, e.g., Fig. B2). Also, beachings identified as occurring on land revealed two situations prior to beaching: (i) an apparently normal drift towards the coast (39 cases, e.g., Fig. B3) and (ii) a sudden change in the direction of the trajectory near the coast (6 cases, e.g., Fig. B4).

Subsequently, the whole dataset of ~ 10 000 identified beaching locations was classified automatically as “in water” or “on land” using an algorithm based on the OpenStreetMap (OSM) land polygons as mentioned in the main text. The automatic classification resulted in an overall percentage of ~53% of beachings in water and ~47% on land, which are very similar to the values obtained in our manual classification of 100 locations. Moreover, a direct comparison of the manual vs. automatic classification of these 100 cases showed that the results were identical except for 3 cases where the locations were actually difficult to determine as being in water or on land, such as in intertidal zones, a few meters from the coast.

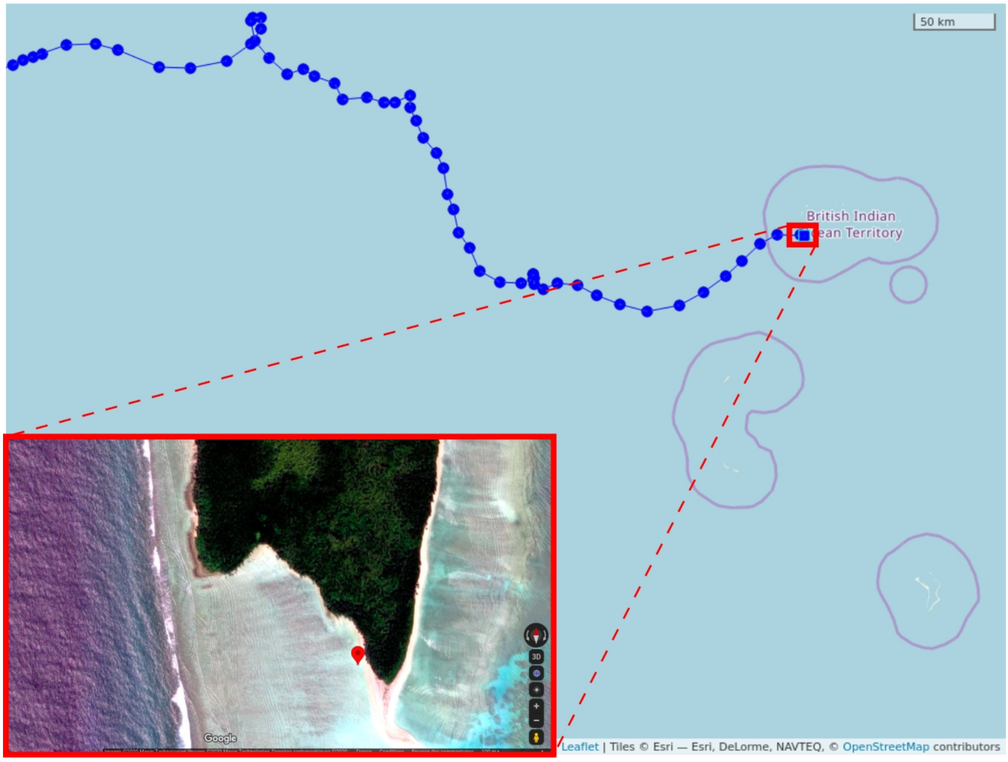

**Figure B1**: The end of the trajectory of buoy n° 50424 (in blue) and a zoom on its identified beaching location (71.7513°,-5.4148°) classified visually as in water.

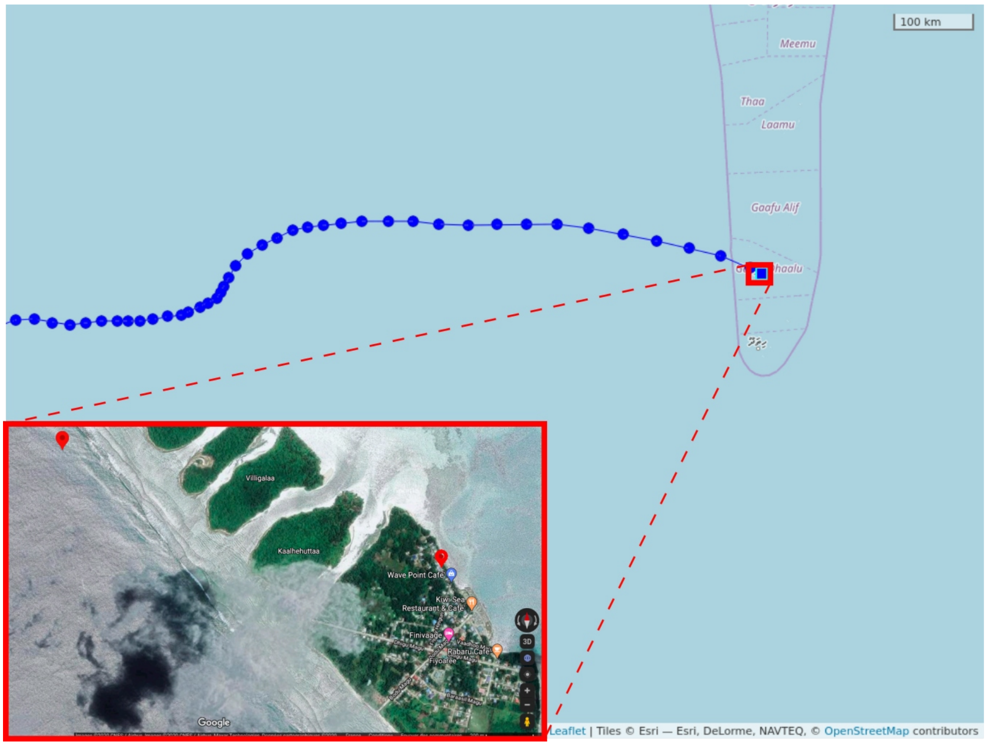

**Figure B2**: The end of the trajectory of buoy n° 9709 (in blue) and a zoom on two identified beaching locations classified visually as in water for the first one (73.1218°,0.2307°) and then on land for the second one (73.1365°,0.226°) 3 days later.

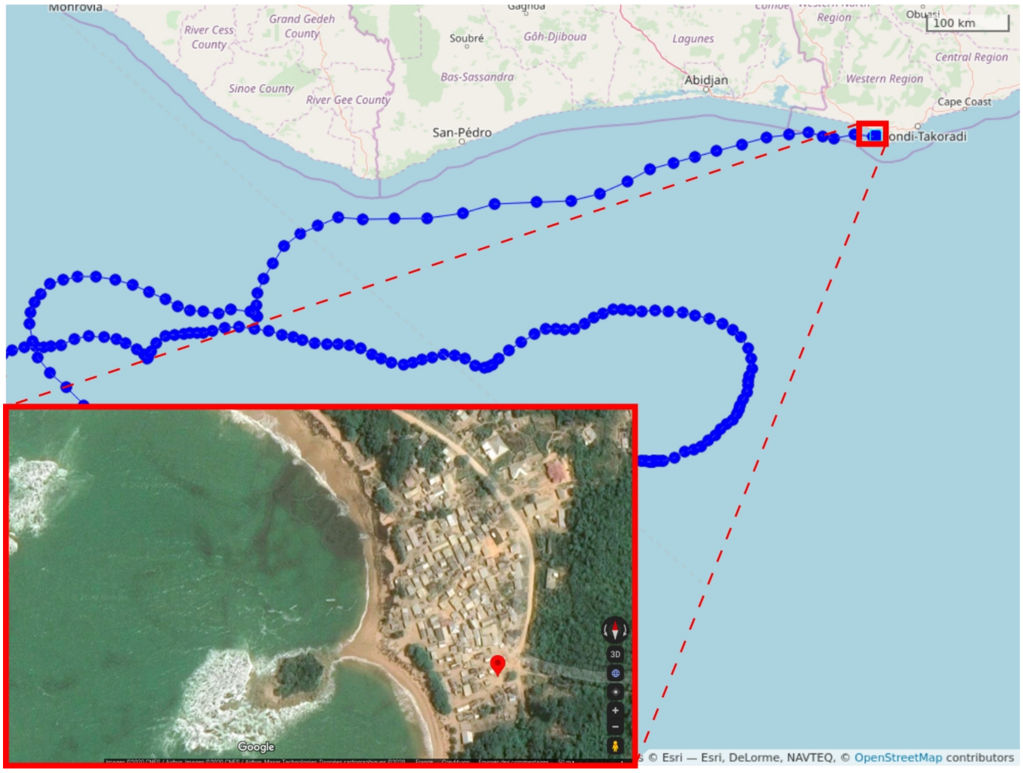

**Figure B3**: The end of the trajectory of buoy n° 30126 (in blue) and a zoom on its identified beaching location (-2.2013°, 4.8303°) classified visually as on land.

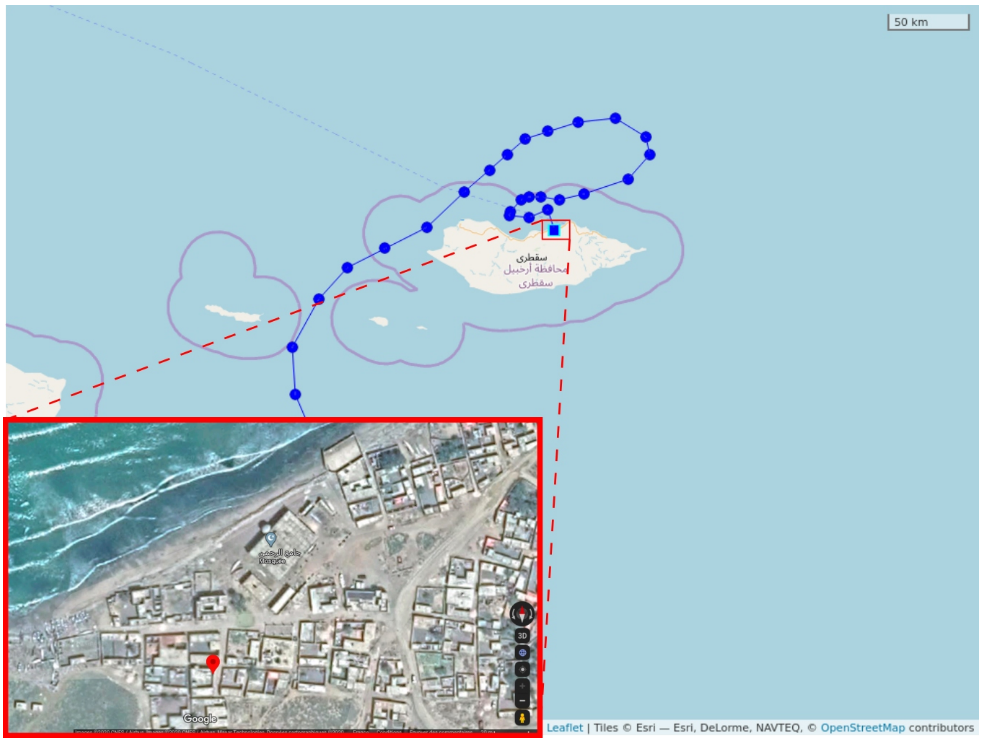

**Figure B4**: The end of the trajectory of buoy n° 15130 (in blue) and a zoom on its identified beaching location (54.016°,12.6522°) classified visually as on land. Note that here there was a sudden change in the trajectory prior to beaching, likely because the buoy was picked up by a boat.

### Appendix C - Additional figures

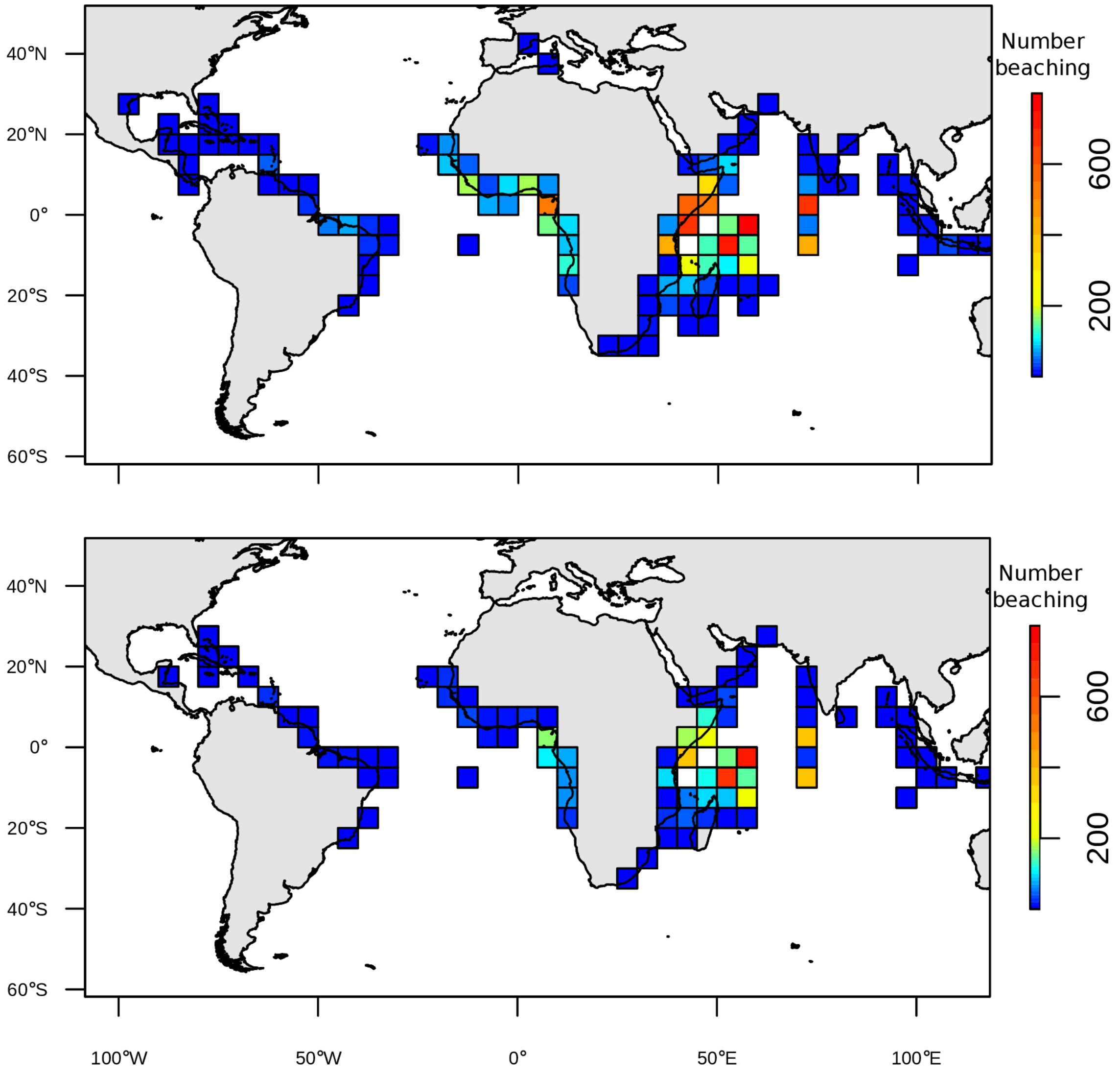

**Figure C1:** The total number of dFADs beached in each 5°x5° grid cell for the period 2008-2017. Redder colors indicate higher numbers of beaching. In (a), all beachings are considered, whereas in (b) only beachings along shore are included. Beachings along shore and recoveries displaced to shore were separated via intersection with OpenStreetMap land polygons. Note that our dFAD trajectory data is incomplete before ~2010, so the absolute number of beachings is likely somewhat higher than values shown in the figure, though differences are likely to be small as the number of dFADs was far lower before 2010 than after 2010.

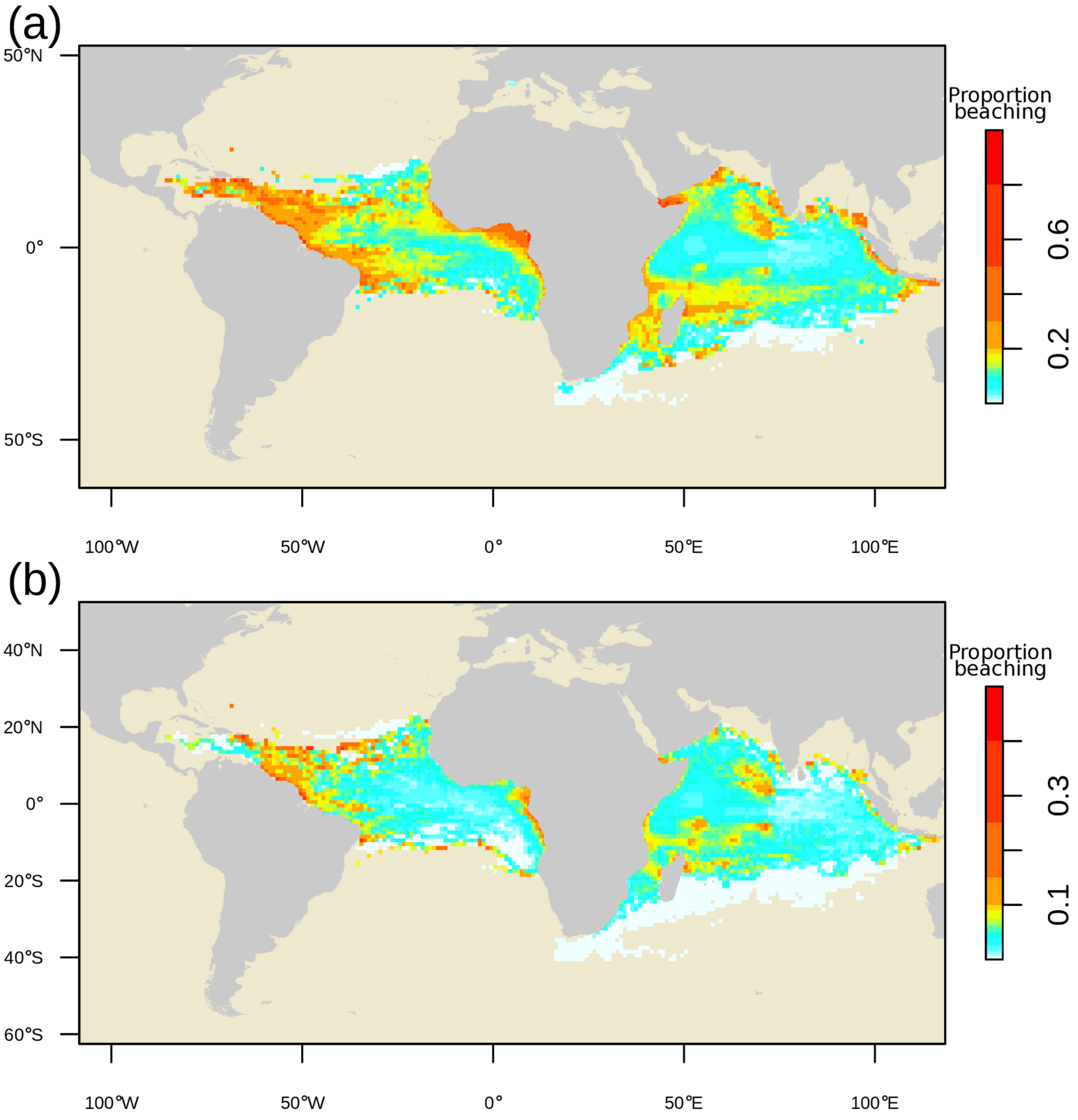

**Figure C2**: Maps of the proportion of dFADs that beached within 12 months after passing through each 1°x1° grid cell over the period 2008-2017. In (a), all beachings are considered, whereas in (b) only beachings along shore are included. The color intervals are unevenly distributed to highlight the low values.

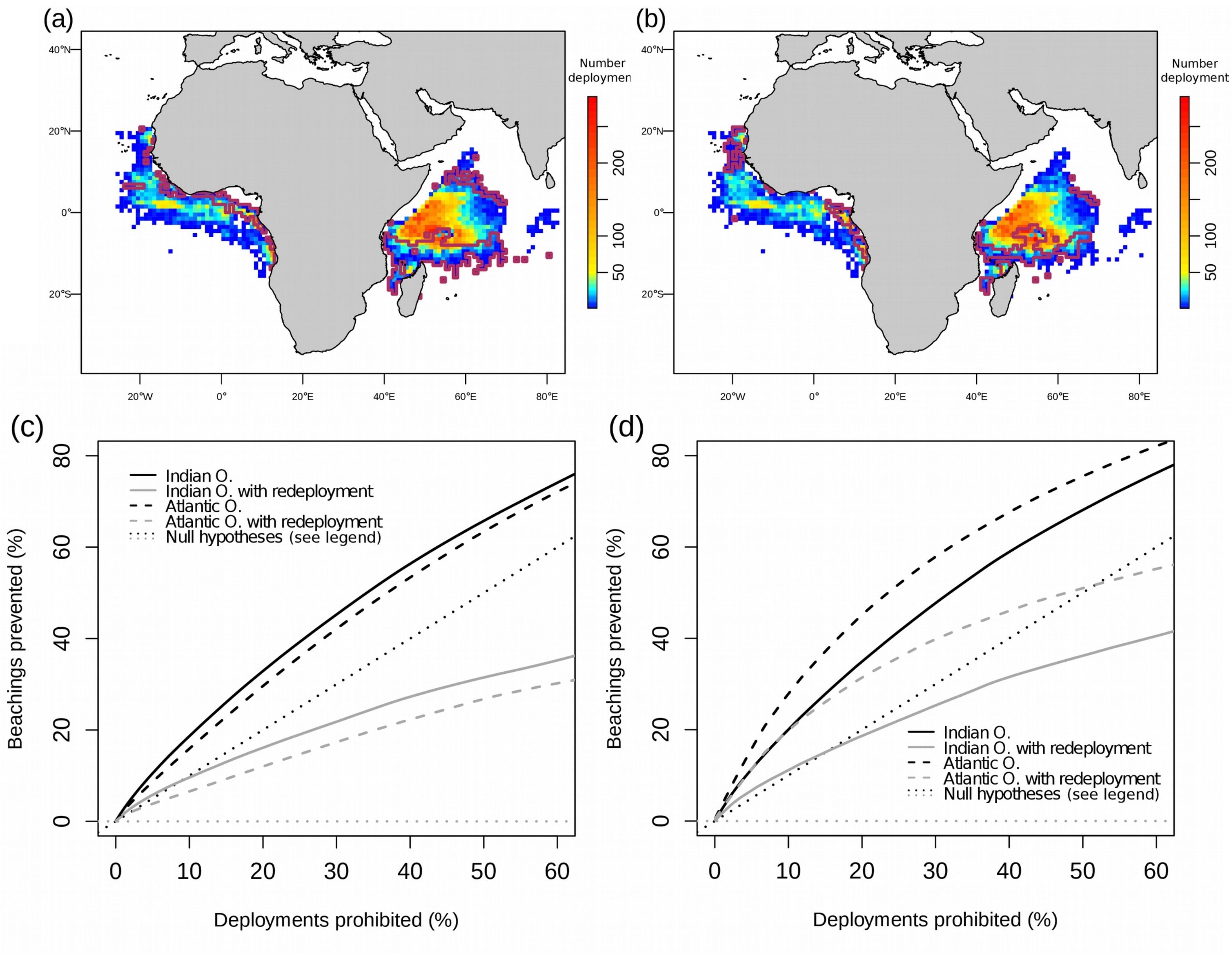

**Figure C3**: Density maps representing the number of dFAD deployments in each 1°x1° cell recorded in logbook data for the period 2013-2017. The purple curves delimit areas representing the 20% of deployments most likely to produce a beaching within 12 months of a dFAD passing through those areas (a-b). Predicted reduction in beaching rate as a function of the amount of area put aside in annual closures to dFAD deployments. Areas are closed from most likely to least likely to produce a beaching within 12 months of deployment (c-d), with area being quantified along the x-axis in terms of the fraction of deployments that occurred in closed areas prior to their closure. In (a-c), all beachings are considered, whereas in (b-d), only beachings  along shore are included.

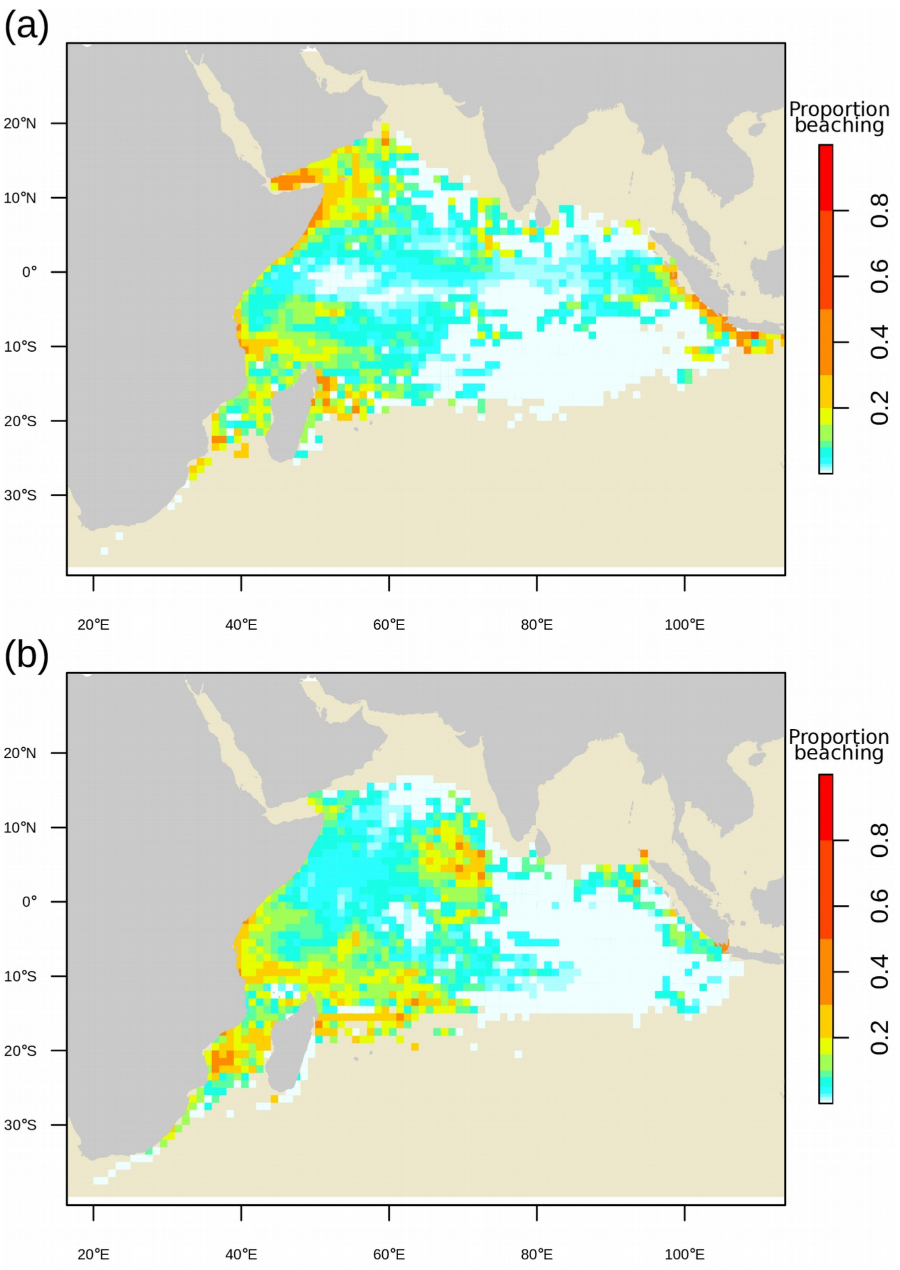

**Figure C4**: Seasonal maps of the proportion of dFADs that beached within 3 months after passing through each 1°x1° grid cell over the period 2008-2017. (a) October-March; (b) April-September. The color intervals are unevenly distributed to highlight the low values.

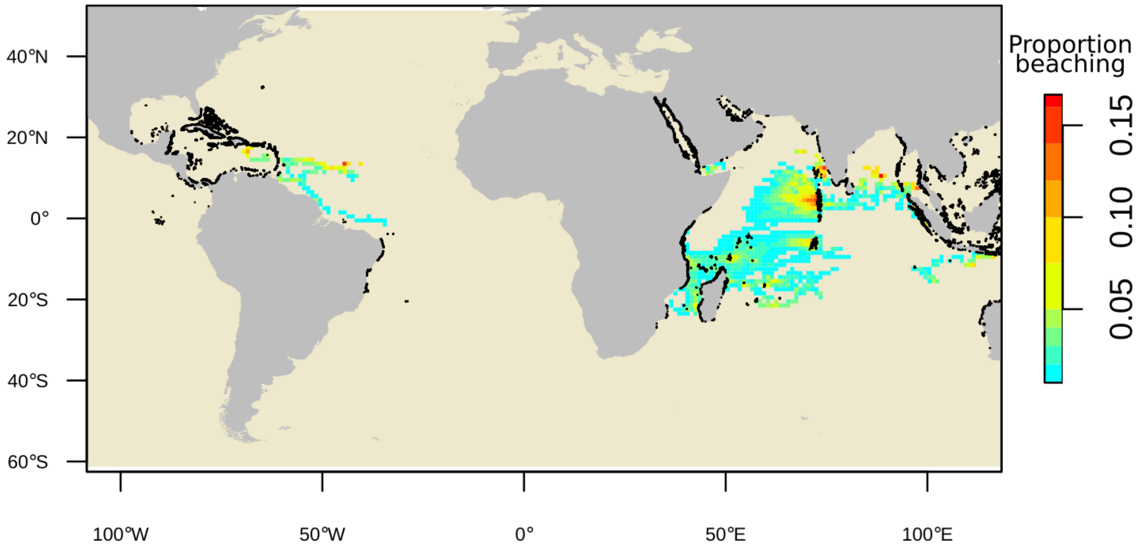

**Figure C5**: Map of the proportion of dFADs that beached **in coral reefs** within 3 months after passing through each 1°x1° grid cell over the period 2008-2017. The color intervals are unevenly distributed to highlight the low values.

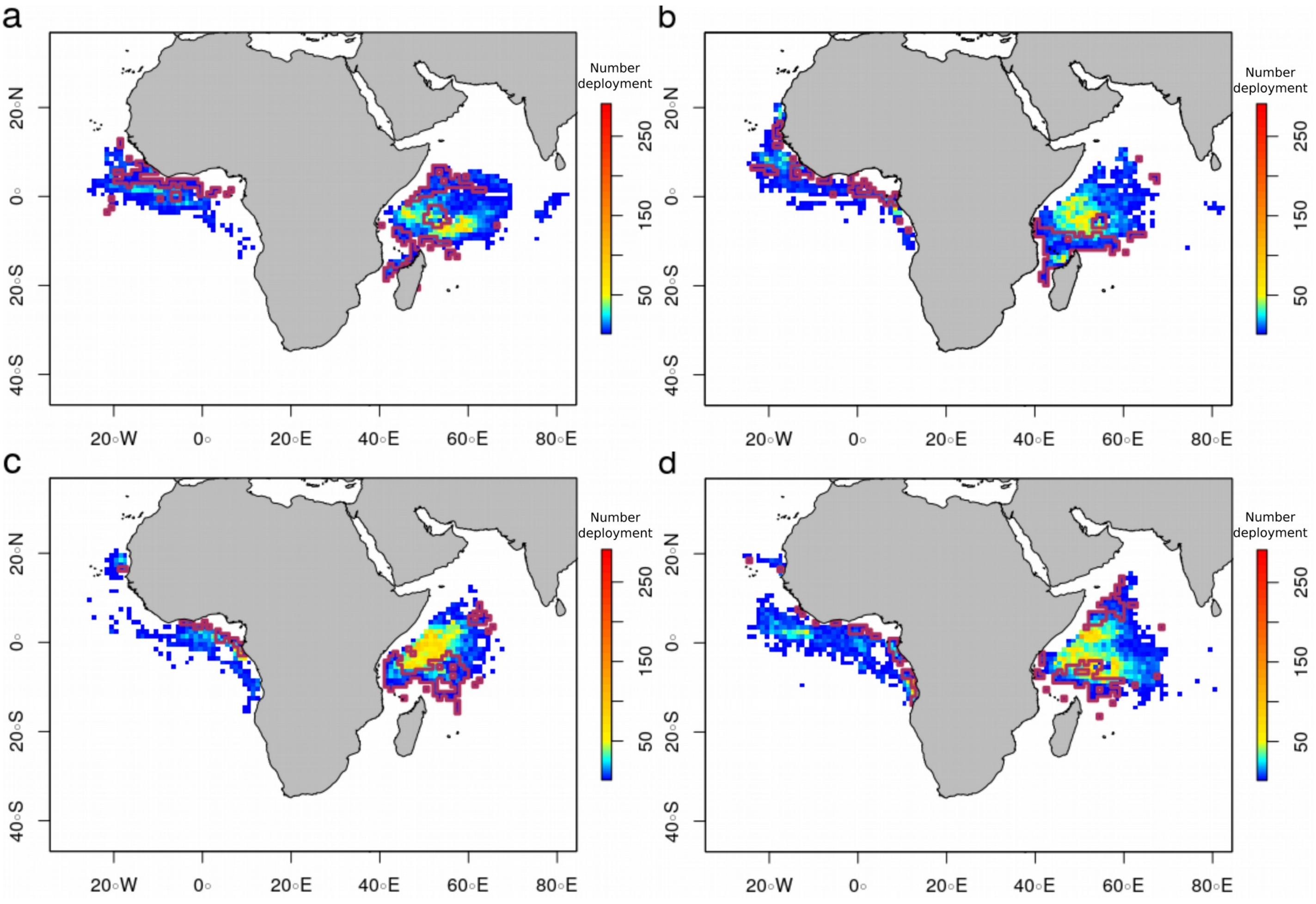

**Figure C6**: Density maps representing the number of dFAD deployments in each 1°x1° cell recorded in logbook data by quarter over the period 2013-2017 (a) Jan-March;(b) Apr-Jun;(c) Jul-Sep;(d) Oct-Dec. The purple curves delimit areas representing the 20% of deployments most likely to produce a beaching within 3 months of a dFAD passing through those areas.

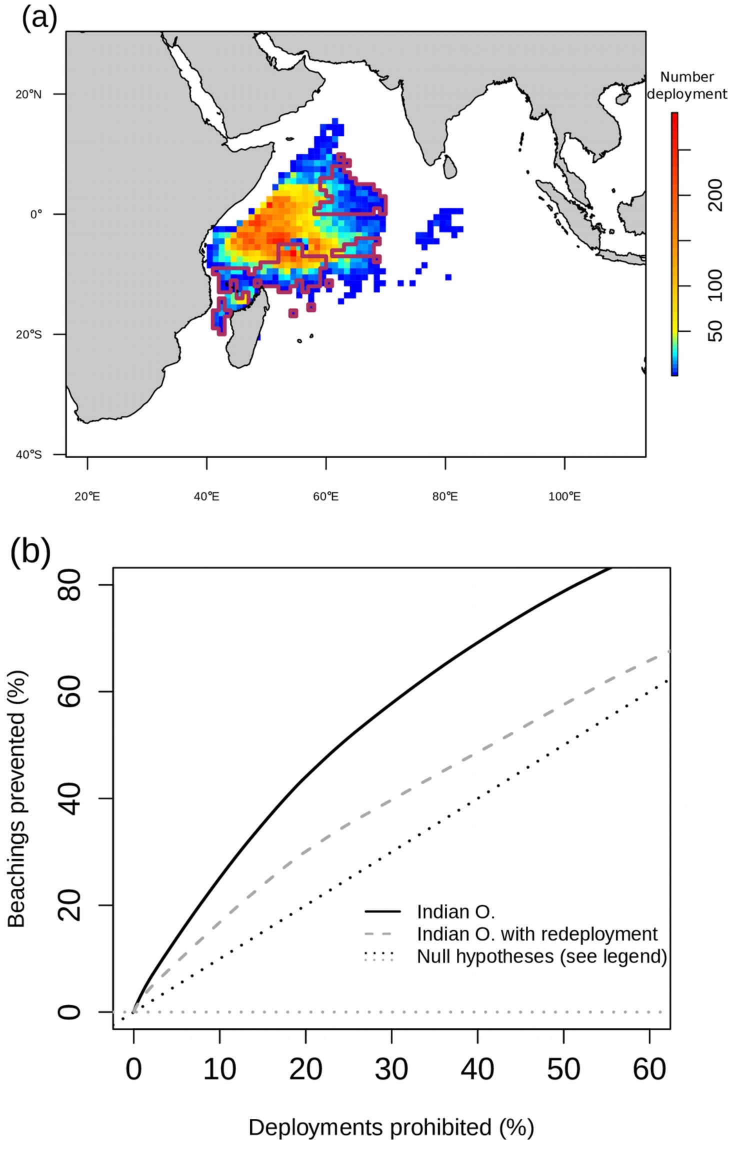

**Figure C7**: (a) Density maps representing the number of dFAD deployments in each 1°x1° cell recorded in logbook data for the period 2013-2017. The purple curves delimit areas representing the 20% of deployments most likely to produce a beaching in coral reefs within 3 months of a dFAD passing through those areas. (b) Predicted reduction in beaching rate as a function of the amount of area put aside in annual closures to dFAD deployments. Areas are closed from most likely to least likely to produce a beaching within 3 months of deployment, with area being quantified along the x-axis in terms of the fraction of deployments that occurred in closed areas prior to their closure.

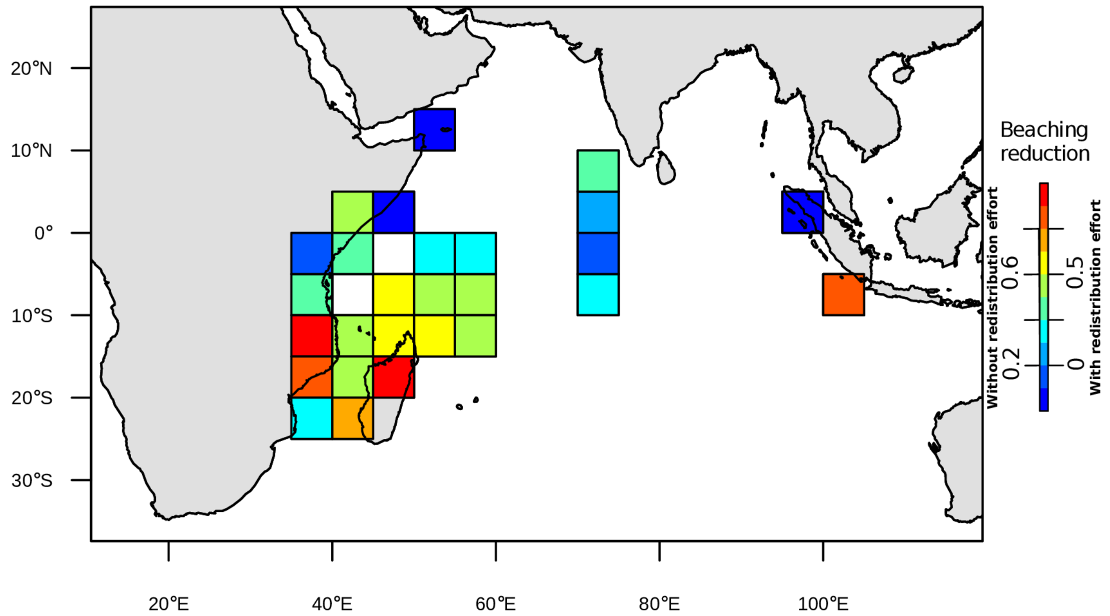

**Figure C8**: Map representing the predicted reduction in beaching when the 20% of dFAD deployments most likely to produce a beaching on coral reefs within 3 months are prohibited (see areas in Fig C7a), without (values on the left of the colorbar) and with (values on the right of the colorbar) dFAD deployment effort redistribution to non-prohibited areas.
